# ANKRD55 interacts with an IFT-B-like complex in microglia

**DOI:** 10.1101/2024.03.18.584613

**Authors:** Jorge Mena, Raquel Tulloch Navarro, Javier Díez-García, Mikel Azkargorta, Ane Aldekoa, Ainhoa Fiat, Nerea Ugidos-Damboriena, Cecilia Lindskog, Olatz Pampliega, Iraide Alloza, Yohei Katoh, Kazuhisha Nakayama, Felix Elortza, Koen Vandenbroeck

**Affiliations:** Inflammation & Biomarkers Group, Biobizkaia Health Research Institute, Barakaldo, Spain; Microscopy Facility, Biobizkaia Health Research Institute, Barakaldo, Spain; Proteomics Platform, Center for Cooperative Research in Biosciences (CIC bioGUNE), Basque Research and Technology Alliance (BRTA), CIBERehd, Bizkaia Science and Technology Park, Derio, Spain; Department of Immunology, Genetics and Pathology, Cancer Precision Medicine Research Unit, Rudbeck Laboratory, Uppsala University, Uppsala, Sweden; Department of Neuroscience, Faculty of Medicine and Nursery, Universidad del País Vasco (UPV/EHU), Leioa, Spain; Laboratory of Glial and Neuronal Autophagy, Achucarro Basque Center for Neurosciences, Leioa, Spain; R&D+I Department, FAES FARMA S.A., Leioa, Spain; Department of Physiology, Faculty of Medicine and Nursery, Universidad del País Vasco (UPV/EHU), Leioa, Spain; Department of Physiological Chemistry, Graduate School of Pharmaceutical Sciences, Kyoto University, Kyoto, Japan; Hiroshima University Genome Editing Innovation Center, 3-10-23 Kagamiyama, Higashi-Hiroshima, Hiroshima, Japan; Department of Biochemistry and Molecular Biology, Faculty of Science, Universidad del País Vasco (UPV/EHU), Leioa, Spain; Ikerbasque, Basque Foundation for Science, Bilbao, Spain

**Keywords:** ANKRD55, intraflagelllar transport, multiple sclerosis, autoimmune, IFT, centrosome, microglia

## Abstract

**Introduction:** SNPs associated with genome-wide risk for multiple sclerosis (MS) modulate expression of ankyrin repeat domain protein 55 (ANKRD55). The function of ANKRD55 is not well understood. A role for ANKRD55 in ciliar transport in multiciliated cells has been reported. To gain deeper insight in how ANKRD55 may modulate neuro-inflammatory parameters, we identified the ANKRD55 interactomes from human neuroblastoma, astrocytic, microglial and monocytic cell lines.

**Methods:** Cell lines were transfected with synthetic ANKRD55 RNA in conjunction with nanoparticles. ANKRD55 interactomes were determined by affinity purification coupled to mass spectrometry (AP-MS) and analyzed bioinformatically. Results were validated and interpreted using confocal immunofluorescence microscopy, RNAseq transcriptomics, and a visible immunoprecipitation assay (VIP).

**Results:** Shared among the interactomes were the 14-3-3 isoforms 14-3-3η and 14-3-3βη. Unique to the microglial interactome were eight proteins belonging to the intraflagellar transport complex B (IFT-B). The IFT-B complex is known to mediate anterograde protein trafficking from the base to the tip of cilia. The dimer IFT46-IFT56 was identified as the minimum entity of IFT-B needed to support interaction with ANKRD55. To verify whether ANKRD55 is a ciliar transport protein, we induced ciliogenesis by serum starvation. Primary ARL13B^+^ cilia could be induced in the astrocytic and neuroblastoma, but not microglial, cell lines. By confocal microscopy, ANKRD55 was not detectable in these cilia but was enriched at the basal body. In the microglial cell line, ANKRD55 and IFT-B components were enriched at the centrosome. In two human primary myeloid cell models, monocyte-derived microglia (MoMG) and monocyte-derived dendritic cells (MoDC), we were able to recapitulate the co-localization of ANKRD55 and IFT81 at the centrosome.

**Discussion:** Our work shows that an ANKRD55 – IFT-B-like complex is assembled in microglial cells. Together with the finding that ANKRD55 was not detected in primary cilia, the results suggest that ANKRD55 is associated with an IFT-B pathway that can operate independent of ciliogenesis.

## 1 Introduction

Ankyrin (ANK) repeat domain 55 protein (*ANKRD55;* Chr5q11.2) is the product of a gene containing intronic genomic variants robustly associated with risk for multiple sclerosis (MS), rheumatoid arthritis (RA), Crohn’s disease and pediatric autoimmune diseases (1–4). The risk alleles of these variants are associated with enhanced expression levels in CD4^+^ T lymphocytes of two genes, *ANKRD55* and its neighbor *IL6ST*. The SNP effect has been co-localized with *cis*-methylation quantitative trait locus (mQTL) effects on a series of CpGs located in and around the *ANKRD55* and *IL6ST* genes, with inverse correlations between *ANKRD55* gene expression and *cis* CpG methylations (5–9). A functional biological link between autoimmunity and the ubiquitously expressed IL-6 family cytokine receptor gp130, coded for by the *IL6ST* gene, and its soluble derivatives (e.g. sgp130), is firmly established (10). The biological function of ANKRD55, however, requires clarification, and its functional relevance to autoimmunity remains to be demonstrated. Be that as it may, expression patterns of ANKRD55 insinuate a participatory role in immune-relevant processes. The gene is naturally transcribed at high levels in peripheral blood CD4^+^ T lymphocytes, but not in CD8^+^ T lymphocytes or any other PBMC subsets, and ANKRD55 protein is enriched in nuclear fractions (9). In monocytes, ANKRD55 expression is gradually induced during 6-day differentiation into immature monocyte-derived dendritic cells (MoDC) in the presence of IL-4 /GM-CSF. This induction is accompanied by increased ANKRD55 immunofluorescence in the cytosol and nucleus with a staining pattern in the latter characteristic of nuclear speckles, and is inhibited by maturation of MoDC with IFN-γ/LPS (11). The main MS and RA risk SNP at the *ANKRD55* locus, rs7731626, apart from modulating *ANKRD55* and *IL6ST* expression in CD4^+^ T cells (7), co-localizes with the strongest *cis*-expression (eQTL) for expression of A*NKRD55*, but not *IL6ST*, in whole blood and spleen (GTEx Analysis Release v8)^1^. In eosinophils, which, like CD4^+^ T cells and basophils, express naturally high levels of ANKRD55 (RNA sequencing of flow sorted immune cells)^2^ (12), TGF-β downregulates transcription of the gene (13). In non-immune cells, ANKRD55 appears biologically relevant as well. Silencing of *ANKRD55* in a human preadipocyte cell line increased significantly both their proliferation rate and lipolysis, but appeared not to affect their differentiation, triglyceride levels or insulin sensitization (14). In mouse hippocampal neuronal cells and mouse microglial cells, ANKRD55 protein is constitutively expressed, and its expression is increased under inflammatory conditions (9). ANKRD55 was recently identified as a specific marker for rare, pediatric, highly malignant CNS neuroblastoma with *FOXR2*-activation (15).

*In silico* structural analysis of the ANKRD55 protein provides a few indications regarding its molecular properties. The structure predicted by the AlphaFold artifical intelligence system (16) (**Figure 1**) reveals a succession of nine 33-residue long ANK repeats located almost entirely in the N-terminal half, followed by a mostly disordered structure extending up to the C-terminus. ANK repeats are known motifs exhibiting a helix−turn−helix conformation, and strings of such tandem repeats are predicted to fold into a single linear solenoid ANK domain. ANK domains are found in hundreds of proteins and are known to facilitate interaction with a diverse range of other proteins with varying sizes and shapes (17–19). For illustration, as shown for the ankyrins (*ANK1/2/3*), which contain 24 ANK repeats, the inner groove spanning the complete ANK repeat solenoid contains multiple semi-independent binding sites capable of engaging different target proteins with very diverse sequences through combinatorial usage of these sites (20, 21). Identification of ANKRD55 interactors could help to uncover specific biological processes that are indicative of its function. The results of several affinity-purification coupled to mass spectrometry (AP-MS) interactome studies of ANKRD55 have been reported. ANKRD55 emerged from the integrated hu.MAP human protein complex map as a component of the ciliar intraflagellar transport (IFT)-B multiprotein complex (22). This complex is known to mediate anterograde protein trafficking from the base to the tip of cilia powered by kinesin-2 motors, and is structurally different from the IFT-A complex that ascertains retrograde protein trafficking via the dynein-2 complex (23). An ANKRD55-GFP fusion protein localized to the cilia of *Xenopus laevis* multi-ciliated epithelial cells, in which it was seen to traffic up and down by means of time-lapse video (22). Morpholino antisense oligo nucleotide knockdown of ANKRD55 led to reduced count and length of cilia, similar to knockdown of its co-IFT-B complex interactor IFT52 (22). Moreover, loss of ANKRD55 was associated with defective vertebrate neural tube closure in *Xenopus* embryos, similar to observations made previously following disruption of IFTs (19, 24, 25). In our recent HEK293 ANKRD55 interactome study, IFT-B components IF74, IFT56 and, less confidently (in two out of three replicates), IFT52, were identified; furthermore, ATP- or nucleotide-binding proteins of diverse functionality with no apparent functional relationships to ciliogenesis were found enriched in the total cell interactome (26).

**FIGURE 1.**
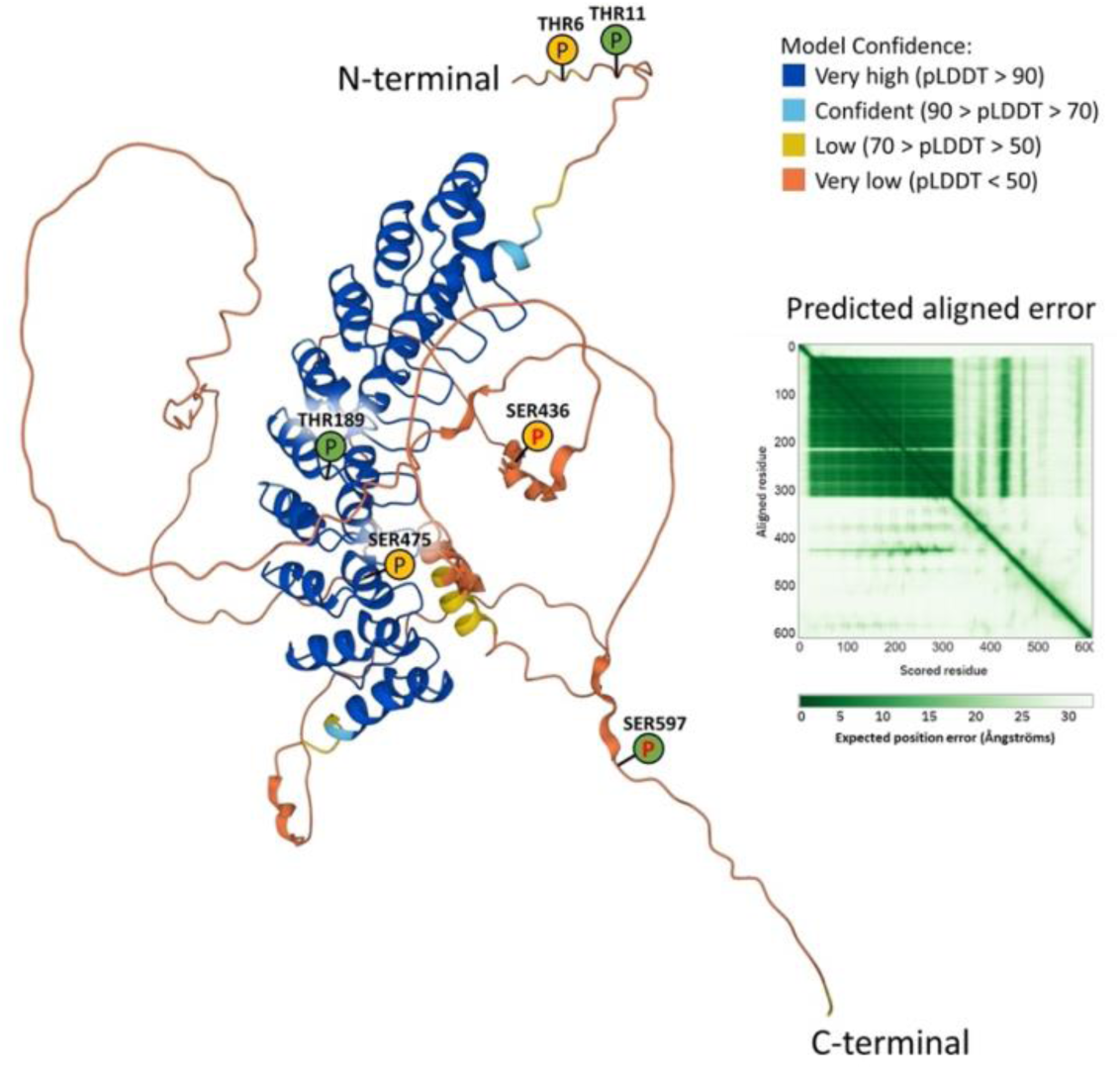
Three-dimensional structure of human ANKRD55 protein (68.4 kDa; Uniprot Q3KP44-1; coded for by full-length transcript ANKRD55-201) predicted by the AlphaFold artificial intelligence system (16). AlphaFold estimates errors using the predicted local distance difference test (pLDDT), which gives a per-residue confidence score from 0 to 100. The model is shown in colors, with high confidence residues colored in blue (including the nine-ANK repeat domain), and low confidence residues in yellow and orange. Predicted aligned error gives a distance error for every pair of residues, indicating if domains are correctly positioned relative to one another. Based on data from Ugidos *et al.* (26), predicted phosphorylation sites (THR11, THR189 and SER597) are indicated with a green circled “P”, and experimentally detected phosphorylation sites (THR6, SER436 and SER475) with a yellow circled P. SER436 and SER597 exhibit the highest scores as likely 14-3-3 binding phosphosite (red symbol P; ref. 26).

ANKRD55 interactomes from two distinct cell lines are available from Bioplex^3^, an unbiased program mapping interactomes of thousands of proteins by AP-MS (27, 28). In Bioplex, the ANKRD55 interactome from HEK293T cells differs markedly from that identified in the colon cancer cell line HCT116 (27, 28). The former displays the IFT-B component proteins IFT52, IFT70A, IFT46, IFT74 and IFT70B as highest probability interacting proteins, while the second is devoid of IFT-B proteins, but contains NAT9, a protein predicted to exhibit N-acetyltransferase activity and PYGM, a glycogen phosphorylase. Shared among both Bioplex interactomes are the oxygen sensor HIF1AN and several 14-3-3-isoforms, conserved regulatory molecules which, through binding to a plethora of functionally diverse signaling proteins, can modulate manifold essential regulatory processes including cell cycle control and apoptosis (29). Cell- or process-specific interactions may thus remodel the ANKRD55 interactome, perhaps reflecting specialized biological pathways in which ANKRD55 participates. IFT-B component proteins are also absent from the ANKRD55 interactome integrated in the human reference interactome (HuRI) map of binary protein interactions (30). In addition to IFT74, various isoforms of the 14-3-3 family of proteins are shared among the Ugidos (26) and Bioplex (28) HEK293(T) ANKRD55 interactomes. Predicted 14-3-3 binding phophosites (26) in the ANKRD55 structure are indicated in **Figure 1**.

In order to expand the interrogable human proteome, to uncover potential cell-type-specified ANKRD55 interactors, and to allow for exploration of processes with potential relevance to neuroinflammation, neurodegeneration and MS, we produced and analyzed by AP-MS the ANKRD55 interactomes from neuroblastoma, astrocytic, microglial and monocytic cell lines. The study confirms 14-3-3 proteins as most shared proteins. Interestingly, an IFTB-like complex was uniquely identified in the microglia cell line. We characterized the structural basis of its interaction with ANKRD55, and studied intracellular co-localization patterns in the absence and presence of serum starvation-induced ciliogenesis. Our data point to a role of ANKRD55 in the life cycle of an IFT-B-like complex associated with the centrosome in the microglia cell line independent of ciliogenesis.

## 2 Materials and Methods

### Cell lines, MoDC and Monocyte-derived Microglia (MoMG)

Immortalized human fetal astrocytes-SV40 (IMhu-A) (Applied Biological Materials, Cat. No. T0280) cells were grown in Roswell Park Memorial Institute (RPMI)-HEPES medium (Sigma-Aldrich, Cat. No. R5886) and cultured in 0.1% collagen I (Sigma-Aldrich, Cat. No. C3867) pre-coated flasks. The immortalized human microglia-SV40 (IMhu-M) cell line (Applied Biological Materials, Cat. No. T0251), HEK293(T) cells and the mouse fibroblast NIH-3T3 cell line (ATCC, Cat. No CRL-1658) were grown in high glucose Dulbecco’s Modified Eagle’s medium (DMEM) (Sigma-Aldrich, Cat. No. D5796). IMhu-M cells were cultivated in 0.1% collagen I pre-coated flasks for improved adherence. IMhu-M cells were cultivated in IL-4/GM-CSF MoDC differentiation medium (Miltenyi Biotec, Cat. No. 130-094-812), when indicated. SH-SY5Y neuroblastoma cell line (Sigma-Aldrich, Cat. No. 94030304) was grown in DMEM/F-12 GlutaMAX™ supplement medium (Thermo Fisher Scientific, Cat. No. 10565-018). THP-1 monocytic cells and Jurkat cells were maintained in RPMI-1640 medium (Sigma-Aldrich, Cat. No. R8758). Alternatively, THP-1 were cultured in MoDC differentiation medium (including IL-4/GM-CSF), as indicated. All cell media were supplemented with 10% heat-inactivated fetal bovine serum (FBS) (Sigma-Aldrich, Cat. No. F9665 or GIBCO, Cat. No. 10270-106), 2 mM L-glutamine (Sigma-Aldrich, Cat. No. G7513), and 1% Penicillin-Streptomycin solution (Sigma-Aldrich, Cat. No. P4458). To generate MoDC, CD14^+^ human monocytes were isolated from fresh peripheral blood mononuclear cells (PBMCs) using CD14 MACS MicroBeads (Miltenyi Biotec, Cat. No. 130-050-201) and cultured in MoDC differentiation medium at a density of 10^6^ cells/ml for 6 days, as described (11). Alternatively, monocytes were cultured in RPMI-1640 in the presence of M-CSF, GM-CSF, NGF-β, CCL2 and IL-34 (cytokines from Peprotech) for 8 days to generate MoMG, following the procedure of Spearman and colleagues (31).

### Synthesis and Transfection of *ANKRD55* mRNA

mRNA was synthesized encompassing the open reading frame of a fusion protein coding for the full-length *ANKRD55* isoform 201 coupled to C-terminal MYC-FLAG tags as provided by a commercial vector (Origene, Cat. No. RC221211). Unmodified synthesized *ANKRD55* mRNA transcript was capped at the 5’ end using wild-type bases CleanCap® AG (TriLink BioTechnologies) to generate a natural Cap 1 structure, that reduces activation of pattern recognition receptors and yields more active mRNA, and the RNA was subsequently polyadenylated (120 A). The mRNA was treated with DNase and phosphatase, purified via silica membrane adsorption, and reconstituted in 1 mM sodium citrate, pH 6.4. Cells were transfected at 60-80% confluency with ANKRD55 MYC-FLAG isoform 201 mRNA using Viromer^®^ mRNA (Lipocalyx, Cat. No. VmR-01LB-00) following manufacturer’s recommendations and incubated for 24 hours. Transfections were performed using 1 µg of *ANKRD55* mRNA per ml of culture media for adherent cells (IMhu-A, IMhu-M and SH-SY5Y), or 2 µg per ml of culture media for suspension cells (THP-1).

### ddPCR and qPCR

Absolute gene transcript quantification was performed by droplet digital PCR (ddPCR). 30 ng of cDNA, ddPCR supermix for probes (no dUTP) (Bio-Rad, Cat. No. 1863023), droplet generation oil (Bio-Rad, Cat. No. 1863005) and specific primers for *ANKRD55* isoform 201 (Bio-Rad) were used to generate droplets in a QX200 Droplet Generator (Bio-Rad). Droplets were then amplified by PCR in a thermal cycler C1000 (Bio-Rad) following manufacturer’s instructions and fluorescence intensity was measured in a QX200 Droplet Reader (Bio-Rad). Data were analyzed using QuantaSoft™ software (Bio-Rad). Quantitative PCR (qPCR) was performed with 5-10 ng cDNA, Fast SYBR^®^ Green Master Mix (Applied Biosystems, Cat. No. 4385612) and specific primer pair best coverage for ANKRD55 isoforms (IDT, Cat. No. Hs.PT.58.27501603201) in a 7500 Fast Real-Time PCR system (Applied Biosystems), following manufacturer’s instructions, and analyzed using the 2^-ΔCt^ method. Samples were assessed in triplicate, no-template controls were included, and *ACTB* (QIAGEN, Cat. No. Hs_ACTB_1_SG) and *GAPDH* (IDT, Cat. No. Hs.PT.39a.22214836) primers were used for normalization. Transcripts quantified by qPCR in the 4 study cell lines were IFT74 (IDT, Cat. No. Hs.PT.58.240526), IFT46 (IDT, Cat. No. Hs.PT.58.26251096), IFT56 (TTC26) (IDT, Cat. No. Hs.PT.58.4747547), IFT81 (IDT, Cat. No. Hs.PT.58.39029302), IFT52 (IDT, Cat. No.Hs.PT.58.19430320), IFT70A (IDT, Cat. No. Hs.PT.58.40272551.g), IFT27 (IDT, Cat. No. Hs.PT.58.45335063), and IFT22 (RABL5) (IDT, Cat. No. Hs.PT.58.15443011).

### Western Blot

Protein cell lysates together with pre-stained protein ladder (Fisher Scientific, Cat. No. BP3603-500), were resolved on 10% SDS-PAGE, transferred to PVDF membranes (Merck Millipore, Cat. No. IPVH00010), blocked with 2% casein (Sigma-Aldrich; Cat. No. C5890) in tris-buffered saline (TBS) for 1 hour at room temperature (RT) and incubated overnight at 4°C with anti-ANKRD55 rabbit polyclonal (1:500; Atlas Antibodies, Cat. No. HPA051049) or anti-FLAG (DDDDK tag) mouse monoclonal (1:800; Proteintech, Cat. No. 66008-2-Ig) primary antibody, followed by incubation with HRP-conjugated donkey anti-rabbit (1:10000; Jackson ImmunoResearch, Cat. No. 711-035-152) or donkey anti-mouse (1:10000; Jackson ImmunoResearch, Cat. No. 711-035-150) secondary antibody for 1 hour at RT. Chemiluminescent signal was detected using enhanced chemiluminescence (ECL) substrate (Bio-Rad, Cat. No. 1705061).

### Flow Cytometry

Untransfected or transfected Jurkat cells were harvested, centrifuged and washed three times with phosphate-buffered saline (PBS). Cells were fixed with 1% paraformaldehyde for 10 minutes at 37°C, permeabilized using ice-cold methanol for 30 minutes on ice, blocked with 3% bovine serum albumin (BSA; Sigma-Aldrich, Cat. No. A9418) for 30 minutes at 37°C, and incubated with FITC conjugated anti-FLAG mouse monoclonal antibody (GenScript, Cat. No. A01632) at a concentration of 2 µg per million cells for 1 hr at 37 C in the dark. Percentage of transfected cells was analyzed in a MACSQuant® flow cytometer (Miltenyi Biotec). CD14^+^ monocytes and inmature MoDC were frozen in fetal bovine serum (FBS) (Sigma-Aldrich, Cat. No. F9665 or GIBCO, Cat. No. 10270-106) with 10% DMSO (Sigma Aldrich ref. SML1661) and subsequently thawed at 37°C for 5 minutes.

The cells were washed with PBS and incubated with LIVE/DEAD Fixable Near-IR Dead Cell Stain reagent (Invitrogen, Cat No. L34975) in order to detect dead cells. Cells were washed with PBS containing 2.5% of bovine serum albumin (BSA; Sigma-Aldrich, Cat. No. A9418) and stained with CD209-PE (Miltenyi, Cat. No. 130-099-707), CD83-APC (Miltenyi, Cat No. 130-098-889) and CD14-FITC (Miltenyi, Cat No. 130-110-576) antibodies for 10 minutes at 4°C in the dark. Percentage of monocytes and immature MoDC cells were analyzed in a MACSQuant® flow cytometer (Miltenyi Biotec). Flow cytometry data were analyzed using FlowJo [version 10.0.7 (TreeStar)].

### Affinity Purification

Untransfected or transfected cells were harvested at 24 h after transfection, centrifuged for 5 minutes at 300 × g and washed three times with PBS prior to protein extraction. Cytoplasmic protein extracts were obtained by incubation of cells with lysis buffer [50 mM NaH2PO4 (Sigma-Aldrich, Cat. No. S8282), 200 mM NaCl, 1% Triton™ X-100 (Sigma-Aldrich, Cat. No. T8787) and 1% complete EDTA-free protease inhibitor cocktail (Roche, Cat. No. 11873580001); pH 8.0] for 45 minutes on ice, followed by centrifugation at 18.000×g for 15 minutes at 4°C and recovery of supernatant.

Protein concentration was measured with Pierce™ BCA Protein Assay Kit (Thermo Fisher Scientific, Cat. No. 23225) and, for each experiment, identical amounts and volumes of protein lysate from either untransfected or transfected cells were added to 20 μl of anti-DYKDDDDK G1 Affinity Resin (GenScript, Cat. No. L00432), previously washed with TBS 0.1% Tween™ 20 (TBS-T), and the mixture was incubated overnight on a rotating wheel at 4°C. Resin was then washed three times with TBS-T, followed by three more washes with TBS on ice, and elution of ANKRD55 complexes was performed with 100 µl of CLB buffer [2 M thiourea (Sigma-Aldrich, Cat. No. T7875), 7 M urea (PanReac AppliChem, Cat. No. A1049), 4% 3-[(3-cholamidopropyl)dimethylammonio]-1-propanesulfonate hydrate (CHAPS; Sigma-Aldrich, Cat. No. 226947)]. All transfections and AP-MS experiments were independently replicated at least three times for each of the four cell lines. Transfection and purification conditions were identical for all cell lines and replicates.

### In-solution Digestion and Mass Spectrometry Analysis

Eluted protein samples were digested using the filter-aided sample preparation (FASP) protocol (32), in which the protein is adsorbed to Amicon^TM^ membranes (Thermo Fisher Scientific) and digestion with trypsin takes place in 50 mM NH4HCO3. Trypsin was added to a trypsin:protein ratio of 1:20, and the mixture was incubated overnight at 37°C, dried out in a RVC2 25 rotational vacuum concentrator (Martin Christ), and resuspended in 0.1% formic acid (FA). Peptides were desalted and resuspended in 0.1% FA using C18 stage tips (Millipore). Samples were analyzed in a timsTOF Pro with PASEF (Bruker Daltonics) coupled online to a nanoELUTE liquid chromatograph (Bruker). 200 ng samples were directly loaded onto the nanoELUTE liquid chromatograph and resolved using 30-minute gradient runs. Database searching was performed using MASCOT 2.2.07 software (Matrix Science) through Proteome Discoverer 1.4 (Thermo Fisher Scientific) against a Uniprot/Swiss-Prot database filled only with entries corresponding to *Homo sapiens*. For protein identification the following parameters were adopted: carbamidomethylation of cysteines (C) as fixed modification, and oxidation of methionines (M) as variable modification, 20 ppm of peptide mass tolerance, 0.5 Da fragment mass tolerance and up to 2 missed cleavage points, peptide charges of +2 and +3. Relative quantification was carried out using a modified spectral counting method named Normalized Spectral Abundance Factor (NSAF) (33). Briefly, protein spectral counts (the sum of all peptide identifications obtained for a certain protein) are corrected by protein length, yielding the Spectral Abundance Factor (SAF) for each protein. These SAF values are further normalized (NSAF) against the sum of all SAF values in a certain sample, and expressed as a % of the total.

### Protein Ranking

Proteins identified in the interactomes were assigned to one out of four categories, defined as follows; Category A, proteins enriched with an NSAF ratio in transfected (T) versus untransfected control (C) cells (NSAF (T) / NSAF (C)) > 2 in at least 2 out of 3 replicates (and absent in the third replicate, if not enriched), and identified with ≥ 2 unique peptides in at least 2 of 3 replicates; Category B, proteins enriched with NSAF(T) / NSAF (C) > 2 in at least 2 out of 3 replicates (and absent in the third replicate if not enriched), and identified with no unique peptide count requirements; Category C, proteins enriched with a NSAF (T) / NSAF (C) ratio > 2 in 2 of 3 replicates that failed to provide such enrichment in the third replicate (present but not enriched), and with no unique peptide count requirements; Category D, proteins that are rescued because, although they failed to provide an enrichment in the 3^rd^ replicate when analysing NSAF, they did when spectral counting is analysed. Within each category (A, B, C, D), proteins were ranked from highest to lowest of NSAF (T) / NSAF (C) ratio averaged over 3 replicates. Proteins absent in untransfected control cells and exclusively present in transfected cells in a single replicate are given a NSAF (T) / NSAF (C) value of 100. Proteins absent in both untransfected control and transfected cells in a single replicate are given a NSAF (T) / NSAF (C) value of 1.

### Immunofluorescence Microscopy and Image Acquisition

IMhu-M, IMhu-A, SH-SY5Y, NIH-3T3 cell lines and MoDC were seeded on coverslips coated with 0.1 mg/ml poly-D-lysine (Sigma-Aldrich, Cat. No. P7886), and MoMG were seeded on coverslips coated with 0.1% collagen type 1 (Sigma-Aldrich, Cat. No. C3867). After cells were cultured, experimental samples were fixed and permeabilized with 100% methanol for 5 minutes at -20°C, and blocked with 1% bovine serum albumin (BSA; Sigma-Aldrich, Cat. No. A9418) in PBS for 30 min. Where indicated, IMhu-M cells were plated on coverslips coated with poly-D-Lysine (Sigma Aldrich, Cat. No. P0899) at a density 1.3x10^5 cells/well and treated with 100 nM bafilomycin A1 (Baf-A1, dissolved in DMSO at 0.16 mM; Sigma Aldrich, Cat. No. SML1661) for 24 h, while control cells were left untreated. Afterwards, cells were fixed with methanol. Coverslips were sequentially incubated with primary antibodies at 4°C overnight, washed in PBS and incubated with corresponding Alexa Fluor-tagged secondary antibodies for 1 h at room temperature (**Supplementary Table 1**). After rinsing in PBS, cell nuclei were stained with 1 μg/ml 4′,6-diamidino-2-phenylindole (DAPI) (Sigma-Aldrich, Cat. No. D9542) diluted in PBS, for 10 min, rinsed, and mounted with Fluoromount-G (Invitrogen, Cat. No. 00-4958-02). For staining of mitochondria, 250 nM MitoTracker^TM^ probe (Thermo Fisher Scientific, Cat. No. M7512) was used. Fluorescence images were acquired using a confocal Zeiss LSM 880 Airyscan microscope mounting an oil-immersion Plan-Apochromat 63x/1.4 objective. Image files were 2048x2048 pixels size achieving a final spatial resolution of 0.065 μm/pixel, and contained four fluorescence channels digitized at 16-bit grayscale. Each of these was recorded in an individual track to avoid bleed-though and maximize photon capture at a GAsP photomultiplier with a 1 Airy pinhole value. For the visualization of fluorophores, the following laser beams and emission ranges were selected: Hoechst, (Ex: 405nm / Em: 425-505nm), Alexa488 (Ex: 488nm / Em: 493-616), Alexa594 (Ex: 561nm / Em: 570-625 nm), and Alexa647 (Ex: 633nm / Em: 638-753 nm). For each experiment, laser power and digital gain were initially adjusted to avoid pixel saturation across different samples.

### Triple Protein Immunofluorescence Co-localization Venn Diagrams

Spatial relationship of immunofluorescence detection of triplets of different proteins was calculated by using a custom-made macro in Fiji software (34). First, individual 16-bit image processing consisted of spatial Gaussian filtering and manual intensity thresholding during visual inspection, for the definition of positive immunodetection selection and subsequent generation of individual ROI series. In the majority of cases, intensity selection excluded the lower third of the whole pixel intensity histogram of the acquired image. ROIs-defined image selection for each wavelength was copied to a black background RGB image to confirm the manual thresholding and to a black background 16-bit image to show overlay delineated areas. Then, total immunofluorescent area was delineated by merging thresholded 16-bit images and selecting pixels that excluded black color.

Individual wavelengths and combinations of emission intersections were defined by using the Image Calculator function in Fiji and again, selecting pixels that excluded black color. Then, area values of each of the perimeters determined (total immunofluorescence, 488 nm emission, 594 nm emission, 647 nm emission, 488-594 nm intersection, 488-647 nm intersection, 594-647 nm intersection and 488-594-647 nm intersection) were measured and expressed as % of the total immunofluorescence. Image montage describing the analysis routine designed in Fiji software is presented in **Supplementary Figure 1**. Graphical representations of the immunofluorescence spatial relationship were created by introducing area values as % in a venn3_circles function and color-coded with Python, GIMP and Adober Illustrator softwares, respectively. Finally, ROI sets from each fluorophore were shown with their corresponding emission color and total area and triple intersection areas were presented as white perimeters or white filled areas, respectively. Triple intersection areas were also shown overlaid on the thresholded colors merged image as pink filled areas.

### RNAseq Transcriptomic Analysis of IMhu-M Cells

RNA samples were isolated using the NucleoSpin RNA extraction kit (Macherey-Nagel, Cat. No. 740955) from mock transfected and ANKRD55-transfected cells at 40 hr after transfection. Two independent experiments were performed. The concentration and the purity of the final RNA solution were checked on the Nanodrop One spectrophotometer (Thermo Fisher Scientific). The isolated RNA was quantified with the Qubit^TM^ 4 fluorometer (Thermo Fisher Scientific) and RNA integrity was assessed with Agilent RNA 6000 nano kit (Aligent, Cat. No. 5067-1511) using an Agilent 2100 bioanalyzer (Agilent). The average A260/280 ratio was 1.9 (range 1.8-2.1), and the average RNA integrity ratio was between 8 and 10. Libraries were constructed with 750 pg of RNA using the NEBNext Single Cell/Low input library preparation kit for Illumina (Cat. No. E6420S). Library concentration and fragment sizes were evaluated with the Agilent High Sensitivity DNA kit (Cat. No. 5067-4626; Agilent). Libraries were sequenced on an Illumina NovaSeq 6000 sequencer (2x150 pb, paired-end) and generated an average of 300 million reads per sample. The FASTQ files were processed for quality control with the FastQC program (https://www.bioinformatics.babraham.ac.uk/projects/fastqc/). Raw paired-end reads were mapped against the *H. sapiens* GRCh38 reference genome using Tophat 2.1.0 tool. Low quality reads were removed using the packages Samtools 1.2 and Picard tools 2.9.0 followed by transcripts assembly and gene identification using Bayesian inference methods with Cufflinks v2.2.2.

### Visible Immunoprecipitation Assay (VIP)

VIP assays were performed as described in detail previously (35). Briefly, expression vectors for ANKRD55 fused to enhanced green fluorescent protein (EGFP) and for combinations of multiple IFT-B components or single proteins fused to mCherry, were co-transfected into HEK293T cells. After 24 h, expression of fluorescent proteins was confirmed by fluorescence microscopy. Next, lysates of the transfected cells were prepared, and processed for immunoprecipitation using GST-tagged anti-GFP nanobody beads. Finally, the beads bearing immunoprecipitates were observed under a fluorescence microscope. In the VIP assay, when two proteins interact with each other, both GFP and mCherry signals are detectable on the beads. On the other hand, when the two proteins do not interact, only GFP signals are observed. To compare the fluorescence intensity, images are acquired under fixed conditions (exposure time and ISO sensitivity of the camera). The materials bound to the beads were subjected to immunoblotting analysis using anti-GFP and anti-mCherry antibodies after image acquisition.

### Bioinformatics Analysis

ANKRD55 interactome proteins were analyzed with the use of Ingenuity Pathway Analysis (IPA) software from QIAGEN (https://digitalinsights.qiagen.com/IPA), and Fisher’s exact test-adjusted *p* values are provided. STRING, a database of known and predicted protein-protein interactions, version 11.5 (https://string-db.org), was used to analyze protein function associated with the combined interactome proteins and with the non-14-3-3 proteins (36). The textmining option was excluded from interaction sources settings. False discovery rate (FDR)-corrected *p* values are provided for STRING analyses. Venn diagram was created using an online tool (http://www.interactivenn.net) (37), and heatmaps were designed with Microsoft Excel software.

## 3 Results

### Expression of Synthetic *ANKRD55* mRNA

Levels of naturally expressed *ANKRD55* full-length isoform 201 mRNA in the study cell lines IMhu-A (astrocytic), IMhu-M (microglia), SH-SY5Y (neuroblastoma) and THP-1 (monocytic) were quantified by ddPCR. Low numbers of transcript copies were observed in THP-1 (0.45 copies / μl) and IMhu-A (1 copy / μl) cells. *ANKRD55* transcripts counts were higher in the other two cell lines, reaching 6.6 copies / μl in IMhu-M and 8.7 copies / μl in SH-SY5Y cells (**Figure 2 A** and **B**).

**FIGURE 2.**
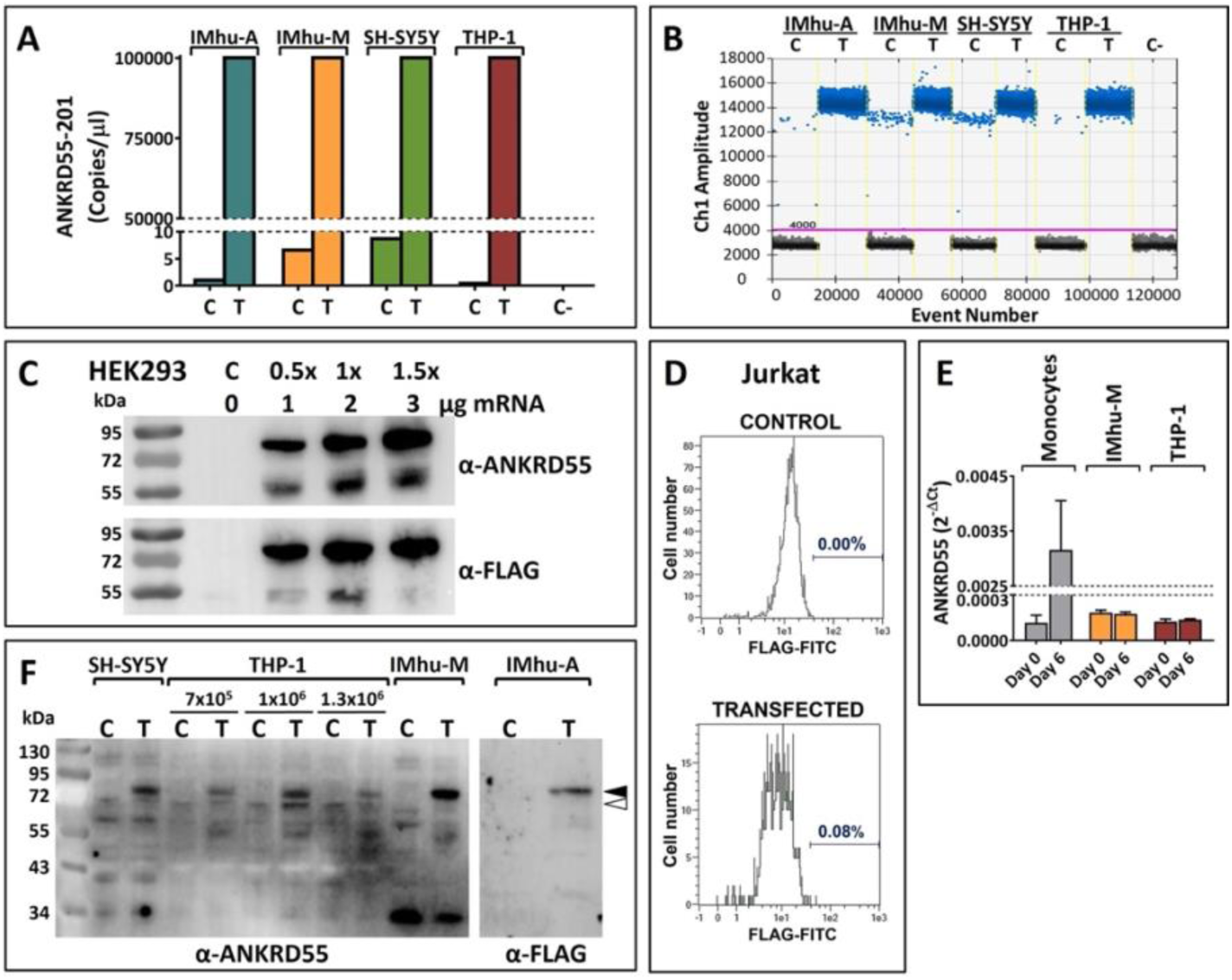
Detection and quantification of native and synthetic mRNA-expressed ANKRD55-Myc-FLAG. (**A,B**) Absolute quantification by digital droplet PCR (ddPCR) of *ANKRD55* mRNA copies in untransfected and transfected immortalized human fetal astrocytes (IMhu-A), immortalized human microglia (IMhu-M), neuroblastoma cells (SH-SY5Y) and monocytic cells (THP-1); representative experiment out of two performed. Number of events are presented in **B**. Positive events (positive fluorescence droplets) are data points in blue above the pink horizontal threshold line, and negative events are those in black below the threshold line. (**C**) Reactivity in Western blot of recombinant ANKRD55 with anti-ANKRD55 and anti-FLAG Abs in lysates from transfected HEK293T cells. (**D**) Lack of detection of ANKRD55 expression in transfected (lower panel) and untransfected (upper panel) Jurkat cells by FLAG-FITC in flow cytometry. (**E**) Induction of ANKRD55 mRNA in primary CD14^+^ monocytes, but not in IMhu-M and THP-1, following 6-day incubation in MoDC differentiation medium containing IL-4/GM-CSF (average of two measurements). (**F**) Western blot analysis of native (white arrowhead) and recombinant ANKRD55 (black arrowhead) in the target cell lines of the study. Equal amounts of cell lysates (10 μg) were loaded in all lanes; numbers above THP-1 lanes indicate density of cells per well tested. C, untransfected control; T, transfected cells.

Transfection scale and expression of synthetic *ANKRD55* mRNA by means of Viromer® polymer nanoparticles was tested in HEK293 cells. 74-kDa protein bands, corresponding to the Mr of full-length protein isoform ANKRD55-201 (69 kDa) with added C-terminal MYC-FLAG tag (**Figure 2 C**), were observed in transfected cells. The transfection method was efficient in all four cell lines, as measured by ddPCR, yielding highly increased *ANKRD55* transcript counts per μl (**Figure 2 A and B**). We also tested whether the T-lymphocytic cell line Jurkat could be transfected with this approach. However, ANKRD55-MYC-FLAG was undetectable in transfected cells via sensitive FLAG-FITC flow cytometry (**Figure 2 D**). Though *ANKRD55* is induced in primary CD14^+^ monocytes during differentiation into immature dendritic cells in the presence of IL-4/GM-CSF (11), this cytokine treatment did not affect native *ANKRD55* expression levels in THP-1 and IMhu-M cell lines (**Figure 2 E**). In western blot of cell lysates of IMhu-A, IMhu-M, SH-SY5Y and THP-1 cell lines, 74-kDa ANKRD55 immunoreactive bands appeared in the transfected cell lines (**Figure 2 F**), at higher intensity than the native 69-kDa protein. Interactome analysis from all cell lines was performed in untreated cells.

### Composition of ANKRD55 Interactomes

Proteins present in ANKRD55 and mock interactomes were identified as described in Materials and Methods. To rank identified proteins, we considered three parameters; enrichment, specificity and abundance of protein detection. The degree of enrichment was represented by the ratio of normalized spectral abundance factor (NSAF) of a protein in interactomes from transfected to untransfected cells. A higher number of unique peptides was considered to enhance reliability of the identified proteins, and overall NSAF value represented peptide abundance in the sample. Four categories (A, B, C, D; see Materials and Methods) were devised, and within each category proteins were ranked according to the average (from highest to lowest) NSAF T/C ratio from 3 independently performed replicates.

The total number of proteins identified in ANKRD55 interactomes per cell line was variable (**Figure 3 A**). The 20 highest ranking proteins per cell line are represented in **Figure 3 B**, and additional identified proteins are provided in **Supplementary Figure 2**. Bioinformatic analysis was performed separately on the combined AB categories (proteins identified following most stringent criteria) or ABCD categories (all identified proteins) proteins. Considering proteins categorized as A or B (AB), two proteins, 14-3-3η (*YWHAH*) and 14-3-3βη (*YWHAB*), were found to be shared among the ANKRD55 interactomes of the four cell lines (**Figure 3 C**). By including also category C and D proteins (ABCD), an additional member of the 14-3-3 isoform family was added to this group, 14-3-3θ (*YWHAQ*).

**FIGURE 3.**
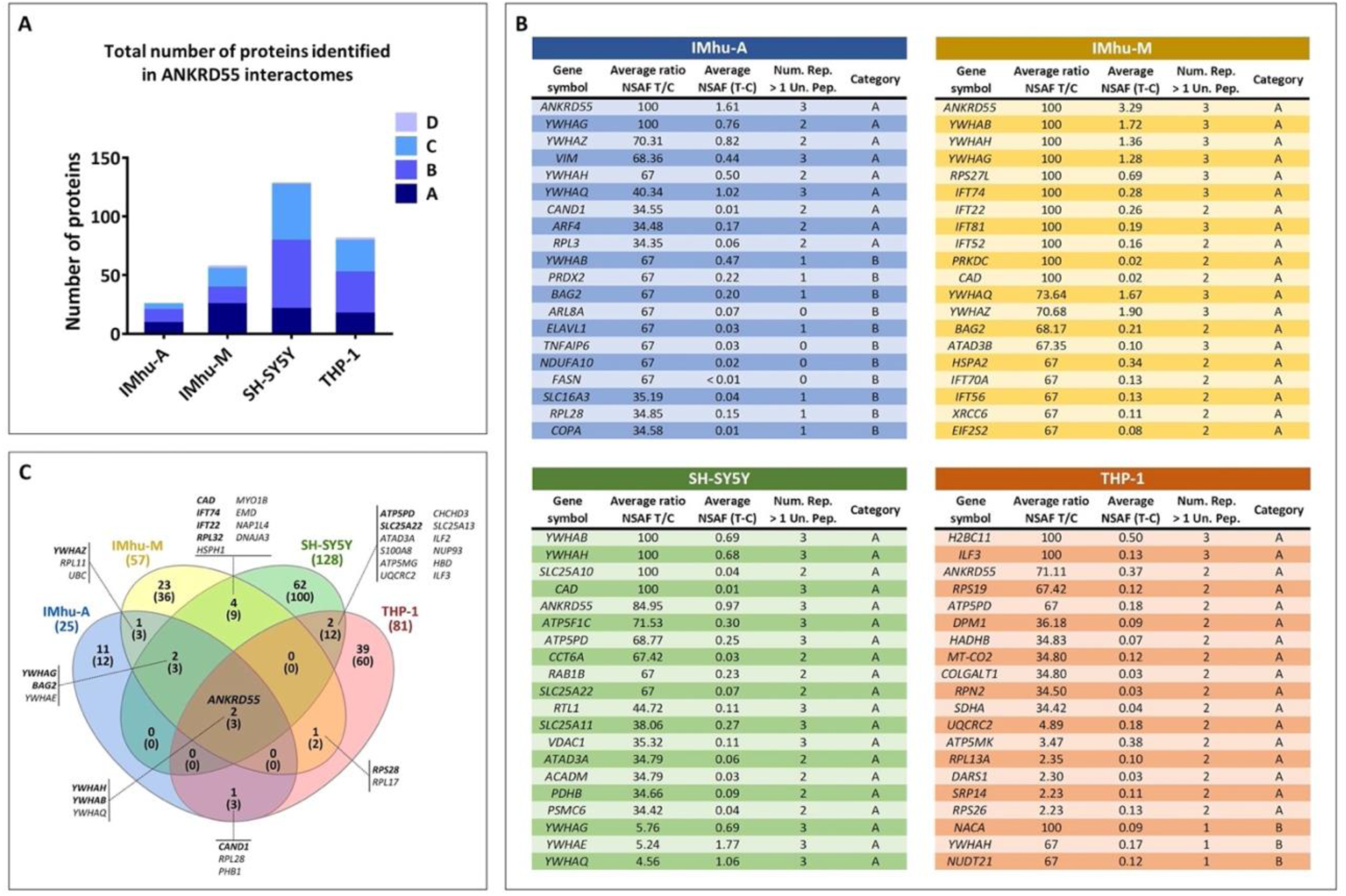
The twenty top-ranked proteins identified by FASP-based AP-MS in ANKRD55 interactomes of IMhu-A, IMhu-M, SH-SY5Y and THP-1 cell lines. (**A**) Number of proteins corresponding to categories A to D (explained in Materials and Methods) identified over three independent replicated experiments for each cell line. (**B**) Top 20 proteins identified in IMhu-A, IMhu-M, SH-SY5Y, and THP-1 interactomes. Proteins are ranked according to the criteria of category A to D, and within each category according to degree of enrichment, i.e. from highest to lowest NSAF (T) / NSAF (C) ratio, averaged over three independent replicates. Proteins absent in the control and exclusively found in transfected cells in a single replicate are given a value of 100 for [Average ratio NSAF (T) / NSAF (C)]. Proteins absent in both control and transfected cells in a single replicate are given a value of 1. Number of replicates in which the protein was identified with more than one unique peptide is indicated (Num. Rep. > 1 Un. Pep.). Protein abundance is provided with NSAF (T – C). (**C**) Venn diagrams of ANKRD55 interactome proteins shared between two, three or all four cell lines over three independent replicates. Number of shared proteins belonging to category AB are shown in bold, with total number of shared proteins (ABCD) indicated below between brackets. Gene symbols of the shared proteins are provided (category AB proteins, italic & bold; category CD proteins, italic).

Three more proteins were shared by the IMhu-A, IMhu-M and SH-SY5Y cell lines, but not by THP-1, i.e. the AB proteins 14-3-3γ (*YWHAG*) and BAG Cochaperone 2 (*BAG2*), as well as the ABCD protein 14-3-3ε (*YWHAE*). No further proteins were shared by any other 3-cell line combinations, though several more proteins were reproduced between pairs of cell lines. The IMhu-M and SH-SY5Y ANKRD55 interactomes had a further four unique AB proteins in common; CAD, a trifunctional multi-domain enzyme involved in the first three steps of pyrimidine biosynthesis; the intraflagellar transport proteins IFT74 and IFT22, and the ribosomal 60S subunit protein RPL32. IMhu-A and IMhu-M shared the sixth 14-3-3 isoform identified as ANKRD55 interactor in this study, 14-3-3ζ/δ (*YWHAZ*). Two mitochondrial membrane proteins, SLC25A22, a glutamate carrier, and ATP5PD, a subunit of mitochondrial ATP synthase, were shared by SH-SY5Y and THP-1 cells. IMhu-M and THP-1 shared 40S ribosomal protein S28 (*RPS28*); and IMhu-A and THP-1 the cullin-associated and neddylation-dissociated protein 1 (*CAND1*).The majority of ANKRD55 interactome proteins were specific to single cell lines (ranging from 50% in IMhu-A to 78% in SH-SY5Y of ABCD proteins).

### Pathways and Functions of ANKRD55 Interactome Proteins

Qiagen IPA software was used to identify canonical pathways associated with shared and cell-specific interactors of ANKRD55. Heatmaps of -log *p* values of associated pathways, and of diseases & biofunctions, are provided for each cell line in **Figure 4** (red panels). Top ranked pathways were shared by the IMhu-A, IMhu-M and SH-SY5Y cell lines. The canonical pathway “Cell Cycle: G2 / M DNA Damage Checkpoint Regulation” emerged with high significance from these three, but with lower significance from THP-1 (*p* values of 1.14×10^-9^, 2.19×10^-12^, 7.89×10^-7^, and 0.006, respectively). Similar observations were made for “HIPPO signaling”, “ERK5 Signaling”, “IGF1 Signaling”, “Protein Kinase A Signaling”, and other pathways. Blue heatmaps representing diversity and relative enrichment of 14-3-3 isoforms in the protein networks associated with each pathway are provided (**Figure 4 A-D**). Different 14-3-3 isoforms were highly enriched in the top ranked pathways of the IMhu-A, IMhu-M and SH-SY5Y cell lines, and non-14-3-3 proteins were rare and not shared among the interactomes, or absent. As an ilustration, the “Cell Cycle: G2 / M DNA Damage Checkpoint Regulation” pathway emerging from category AB proteins refers to the presence of five distinct 14-3-3 isoforms and no other proteins in IMhu-A and SH-SY5Y, to six 14-3-3 isoforms in addition to PRKDC in IMhu-M, and to two 14-3-3 isoforms as only associated network proteins in THP-1 interactomes.

**FIGURE 4.**
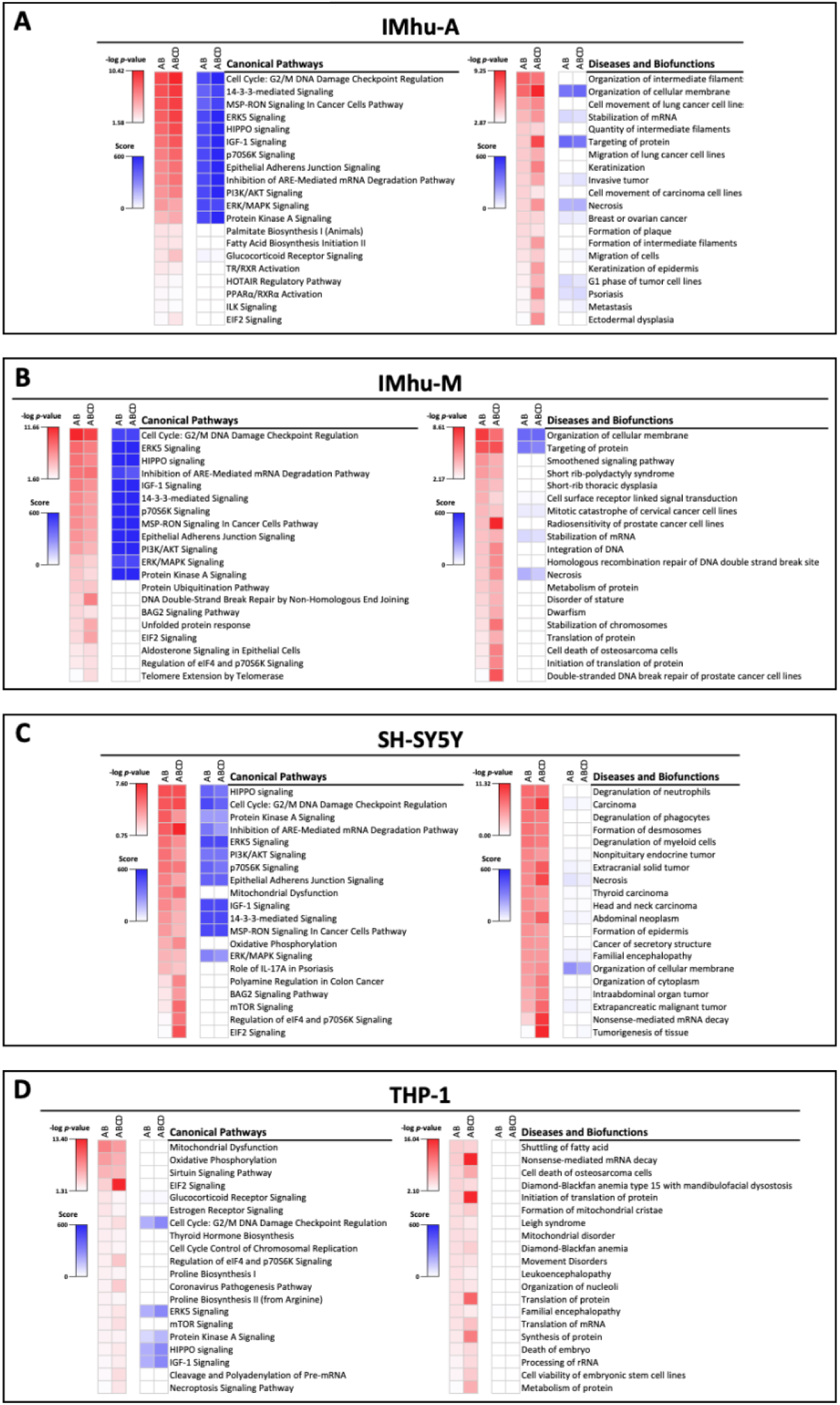
Ingenuity Pathway (Qiagen IPA software) analysis of ANKRD55 interactomes from (**A**) IMhu-A, (**B**) IMhu-M, (**C**) SH-SY5Y and (**D**) THP-1 cells. Red heatmaps provide significance (-log *p* value) of canonical pathways, and of diseases and biofunctions detected in category AB and ABCD interactomes, ranked following the significance levels (high to low) found in the category AB proteins. Blue heatmaps represent the diversity and relative enrichment of 14-3-3 isoforms in the identified protein network assigned to canonical pathways or biofunctions in each cell line. The score of the blue heatmaps was calculated by multiplying the number of distinct 14-3-3 isoform proteins present in the pathway or biofunction with the % of 14-3-3 proteins in the associated protein network. The maximum score of 600 corresponds to the presence of six diverse 14-3-3 isoforms and no other proteins, and is seen in 9 of the 12 top ranked pathways in IMhu-M category AB and ABCD proteins.

The 7^th^ known 14-3-3 isoform, sigma or 14-3-3σ (*SFN*) was not identified in any interactome in this study. RNAseq transcriptomic analysis was performed on the mock-and ANKRD55-transfected IMhu-M cells which showed that the *SFN* gene was not transcribed in either case. We used the data from the RNAseq analysis to assesss how individual transcript read counts relate to associated protein abundancy in the IMhu-M interactome. Figure 5 A shows that RNAseq mRNA read counts of ANKRD55 interactome proteins were positively correlated with their normalized interactome NSAF values (Pearson *r* = 0.397, *p* = 0.002). However, the correlation was entirely driven by the six 14-3-3 isoforms (Figure 5 **B - C**; Pearson *r* = 0.94; *p* = 0.005), the most abundant proteins in the interactome (highest NSAF T – NSAF C values, Figure 5 A), and it trended towards negative in their absence (Pearson *r* = -0.253, *p* = 0.076; Figure 5 B). 14-3-3 proteins may bind to ANKRD55 via its predicted 14-3-3-binding phophosites (Figure 1), though phospho-independent 14-3-3 protein – target protein interactions are known to occur (38). The data show that ANKRD55 is capable of binding all six 14-3-3 isoforms available by RNAseq in the IMhu-M cell line, in proportionality to their respective cellular protein levels estimated by RNAseq.

**FIGURE 5.**
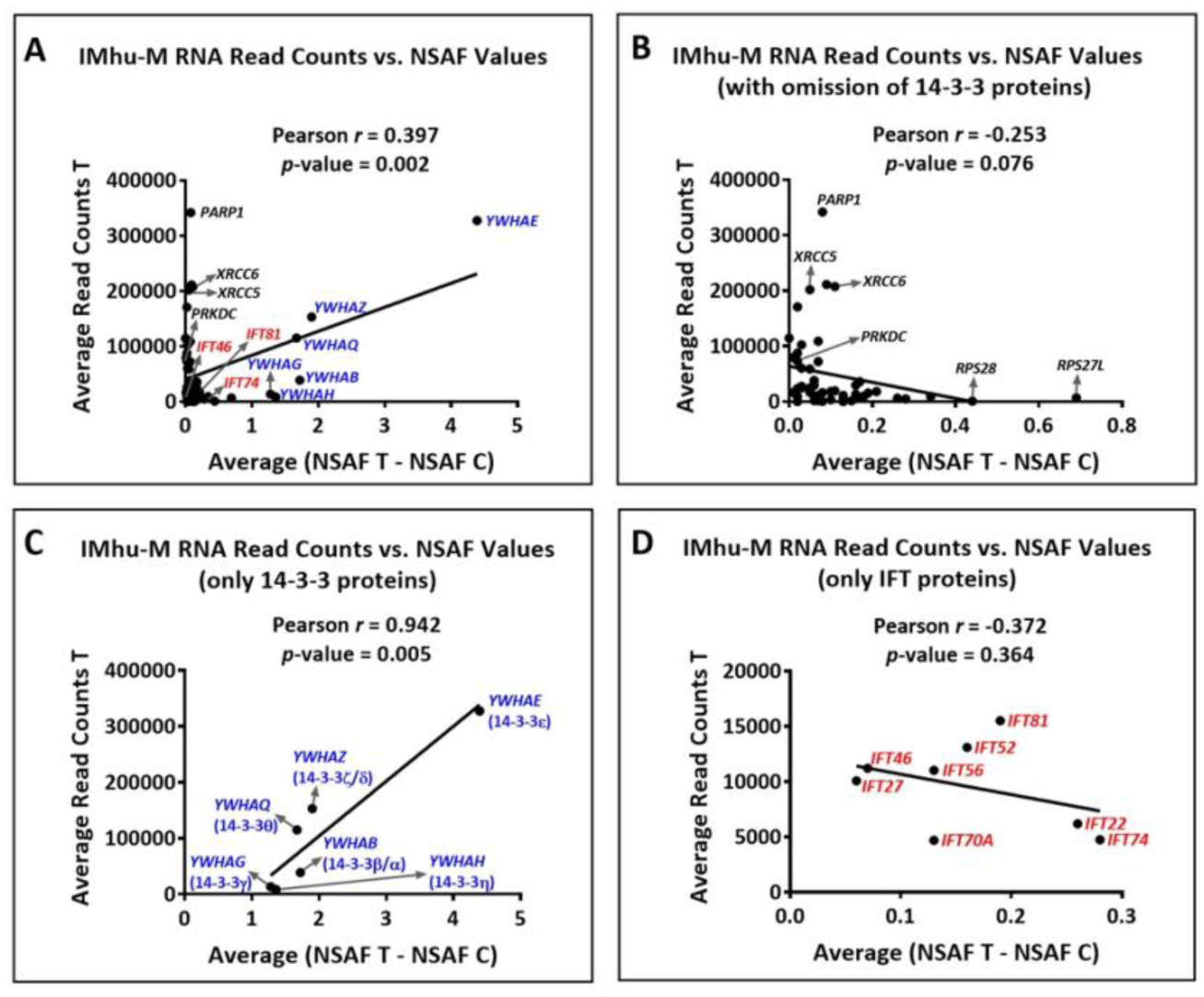
Pearson correlation analysis between cellular RNAseq transcript read counts of proteins identified by AP-MS in the IMhu-M ANKRD55 interactome, and their respective normalized spectral abundance factor (NSAF) values in the interactome [difference of NSAF between transfected (NSAF – T) and untransfected (NSAF – C) values] (average of two and three independent replicates, respectively). The analysis was performed including **(A)** all proteins identified in the interactome, **(B)** excluding 14-3-3 isoforms from total proteins, **(C)** considering only 14-3-3 isoforms or, **(D)** considering only IFT proteins. Pearson correlation coefficients (*r*) and *p*-values are shown. 14-3-3 isoforms are indicated in blue and IFT proteins in red. For reference, some other interactome proteins with high average read count are indicated, specifically the members of the DNA repair complex PRKDC-PARP1-XRCC5-XRCC6 complex (black).

Canonical pathways from which 14-3-3 proteins were absent, had lower significance levels, and were less frequently shared among cell lines (Figure 4). One of these, “Mitochondrial Dysfunction”, the top THP-1 canonical pathway (Figure 4 D) (*p* value for AB proteins: 4.33×10^-9^; protein network: ATP5MG, ATP5PD, CPT1A, NDUFA4, NDUFB4, SDHA, UQCRC2, VDAC3) was identified also in SH-SY5Y cells (Figure 4 C) (*p* value for AB proteins: 2.75×10^-5^; protein network: ATP5F1C, ATP5PB, ATP5PD, CAT, NDUFA13, VDAC1). Only ATP5PD was shared between both networks, but additional discrete subunits of the mitochondrial ATP synthase complex (39) were identified, and different mitochondrial NADH-Ubiquinone Oxidoreductase respiratory chain complex proteins (NDUF) and Voltage Dependent Anion Channels (VDAC) isoforms were also present in both interactomes. Functional enrichment analysis using STRING database was performed following removal of all 14-3-3 isoform proteins from the interactomes, so as to search for patterns in the remaining protein sets. Uniprot Keyword “Acetylation” was the most enriched term in the combined interactomes from the four cell lines (Figure 6 A; FDR *p* = 2.71×10^-32^ in AB and 1.2×10^-53^ in ABCD protein categories), as well as in the individual SH-SY5Y and THP-1 interactomes (Figure 6 B). “Extracellular exosomes” and “RNA binding” were weakly associated with the individual interactomes of three cell lines (Figure 6 **B;** AB protein FDR *p* < 4.7×10^-02^). Compartment terms organelle / mitochondrial in combination with envelope / (inner) membrane were significantly enriched only in the SH-SY5Y and THP-1 interactomes (AB protein FDR *p* < 7.5×10^-3^ and < 1.36×10^-5^, respectively), in accordance with the IPA analysis pointing to association of ANKRD55 with mitochondrial function in these cells.

**FIGURE 6.**
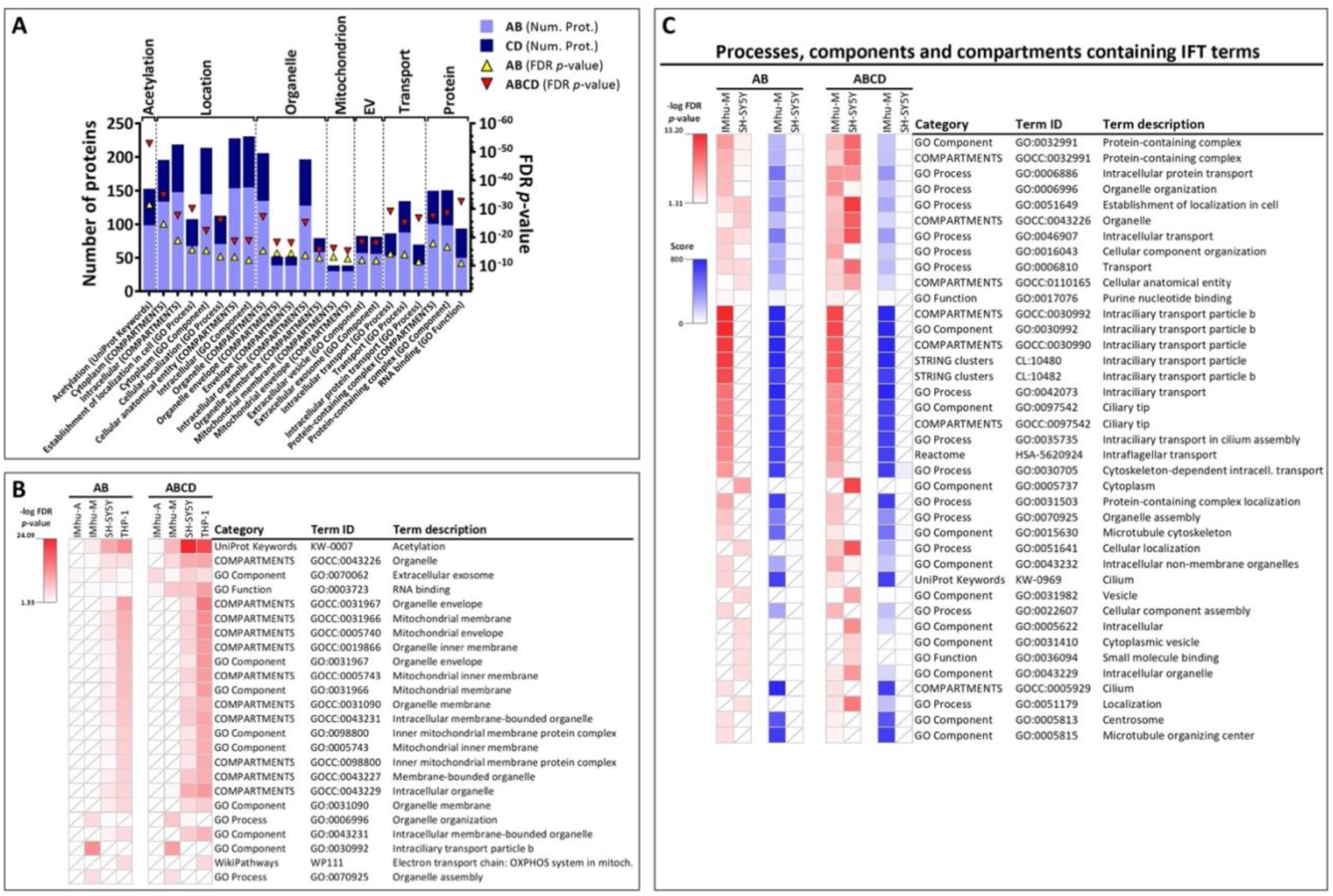
STRING database analysis of 14-3-3-free ANKRD55 interactomes. (**A**) The total number of proteins present in each STRING term from analysis of the combined four-cell line interactomes, omitting all 14-3-3 isoforms, is indicated with bars for category AB proteins (light blue) and category CD proteins (dark blue). Significance associated to each STRING term, expressed as an FDR *p*-value, is represented with yellow triangles for category AB proteins and red triangles for category ABCD proteins. Terms obtained from the STRING analysis were grouped according to similarity in cell function and location. (**B**) Heatmap from STRING analysis performed on the separate category AB and ABCD ANKRD55 interactomes of IMhu-A, IMhu-M, SH-SY5Y and THP-1 cell lines, excluding all 14-3-3 protein isoforms. Significance values are provided as colour-coded -log FDR *p*-values. Cells with a diagonal line indicate an absence of a STRING term. (**C**) STRING analysis terms containing IFT proteins from IMhu-M (IFT22, IFT74, IFT81, IFT52, IFT56, IFT46, IFT70A, IFT27) and/or SH-SY5Y (IFT22, IFT74) interactomes. Statistical significance (expressed as -log FDR *p*-value) of STRING terms associated with category AB and ABCD protein interactomes is shown as red heatmaps. Blue heatmaps show the diversity and relative abundance of IFT proteins in each STRING term. A score, calculated by multiplying the number of distinct IFT proteins detected with the % of IFTs in the STRING terms, was assigned to each term. The maximum score of 800 corresponds to presence of eight different IFTs and no other proteins. Absence of a STRING term in a cell line is represented with a diagonal line in the corresponding cell.

Interestingly, GO Component “Intraciliary transport particle b” (GO:0030992) emerged as the most enriched STRING term of all cell lines (AB proteins), but was uniquely associated with the IMhu-M ANKRD55 interactome (Category AB FDR *p* = 6.7×10^-14^) (Figure 6 B). The interactome protein network of this pathway contained 8 category AB proteins, IFT74, IFT22, IFT81, IFT52, IFT70A, IFT56, IFT46, and IFT27 (Figure 3, **Supplementary Figure 2**), and adding category CD proteins to the analysis did not reinforce the enrichment (Figure 6 B). In the IPA IMhu-M ANKRD55 interactome analysis (Figure 4 B), diseases or functions annotated to some members of this group of IFT proteins included: the smoothened branch of Hedgehog pathway for signaling across the membrane (IFT27, IFT46, IFT52, IFT81, IFT56), cell surface receptor linked signal transduction (IFT27, IFT46, IFT52, IFT81, IFT56, in addition to UBC, YWHAZ), the skeletal ciliopathies short-rib polydactyly syndrome & thoracic dysplasia (IFT52, IFT74, IFT81), disorder of stature (IFT52, IFT74, IFT81, in addition to DSP, XRCC6), and dwarfism (IFT52, IFT74, IFT81, in addition to XRCC6). Although two of these proteins, IFT22 and IFT74, were also identified in the SH-SY5Y interactome, analysis of IFT-containing terms in both these interactomes by STRING database, did not uncover shared informative categories beyond GO terms reflecting moderately enriched basic processes such as (intracellular) transport and establishment of localization in cell (Figure 6 C). No correlation was found between IMhu-M RNAseq read counts of the identified IFTs and their individual normalized NSAF values in the ANKRD55 interactome (Figure 5 D; Pearson *r* = -0.372 ; *p* = 0.36). In contrast to the 14-3-3 proteins (Average NSAF (T) – NSAF (C) = 1.28 – 4.39), IFT proteins interacted with ANKRD55 over a much lower, and more narrow normalized spectral abundancy interval (Average NSAF (T) – NSAF (C) = 0.06 – 0.28).

### ANKRD55 Interacts with the IFT-B Heterodimer IFT46 – IFT56

Given this data, and the earlier data in (22) revealing ANKRD55 as a member of the IFT-B complex, we sought to study the molecular basis of the interaction between ANKRD55 and the identified IFT-B components. The known IFT-B holocomplex mediating ciliary protein trafficking is well-characterized. It is structurally divided into two subcomplexes; the core subcomplex (also referred to as the IFT-B1 subcomplex) composed of 10 subunits (IFT22/IFT25/IFT27/IFT46/IFT52/IFT56/IFT70/IFT74/IFT81/IFT88), and the peripheral subcomplex (the IFT-B2 subcomplex) composed of six subunits (IFT20/IFT38/IFT54/IFT57/IFT80/IFT172) (23). The IFT-B1 (core) subcomplex itself consists of core-1 and core-2 subcomplexes each containing 5 distinct IFTs. As shown in Figure 7 A, the 8 IFT-B subunits identified in the IMhu-M ANKRD55 interactome are scattered over the known IFT-B core-1 and core-2 subcomplexes, and are absent from the peripheral IFT-B2 subcomplex.

**FIGURE 7.**
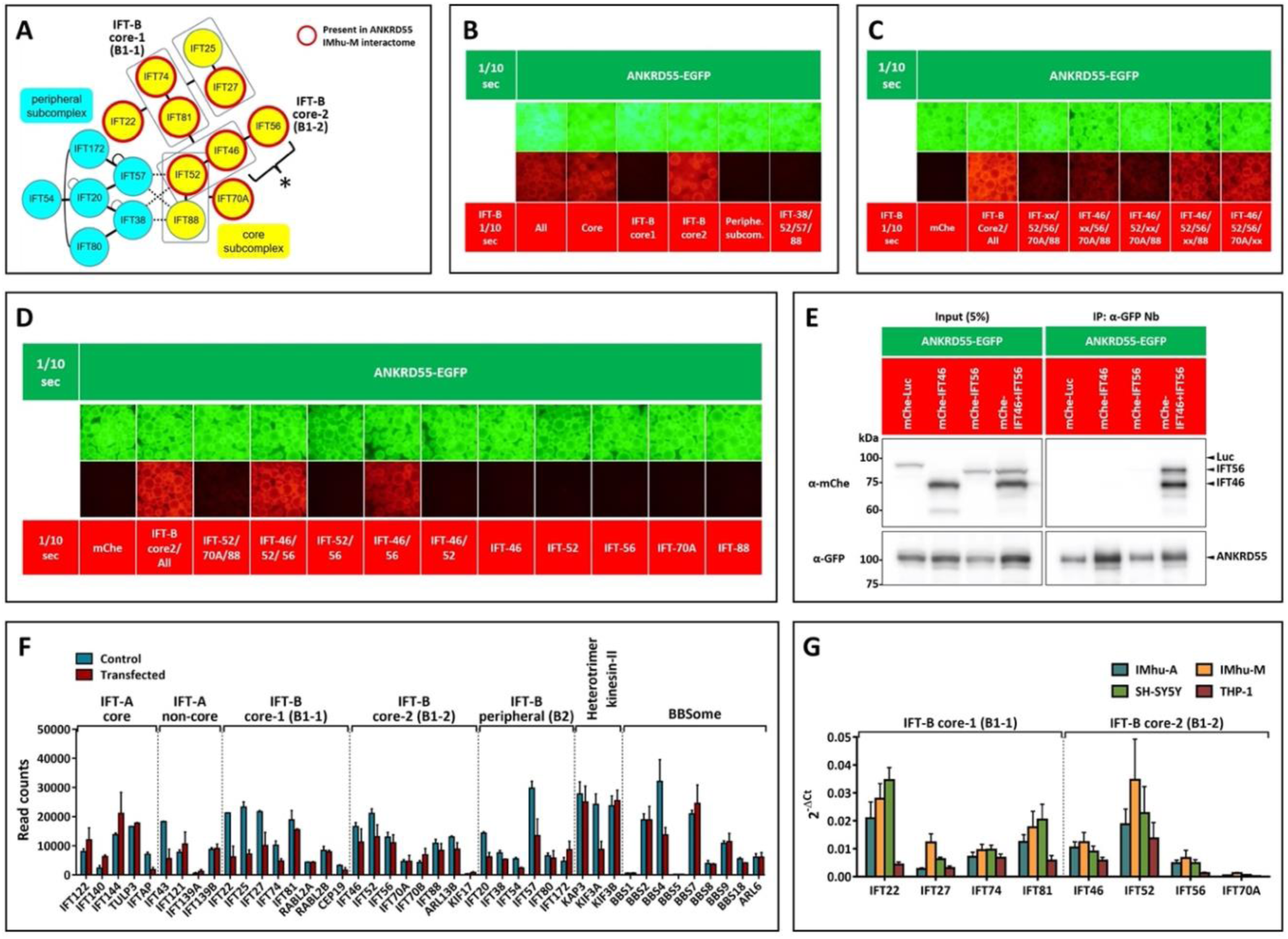
VIP assay analysis of the interaction of IFT-B components with ANKRD55. (**A**) Architecture of the IFT-B complex. Subunits of the core and peripheral subcomplexes are colored yellow and light blue, respectively. Broken lines indicate composite interactions involving IFT38, IFT52, IFT57, and IFT88 (23). Proteins detected in the IMhu-M ANKRD55 interactome are circled in red. * indicates the binding site of ANKRD55 determined in **B-E**. (**B, C, D, E**) VIP assay to identify ANKRD55-binding IFT-B components. HEK293T cells cultured in 6-well plates were transfected with expression vectors for EGFP-ANKRD55 and for combinations of multiple IFT-B components or single proteins fused to mCherry. 24 h after transfection, lysates were prepared from the cells and precipitated with GST-tagged anti-GFP nanobody prebound to glutathione-Sepharose beads and then processed for the VIP assay (**B, C, D**) or immunoblotting analysis using antibodies against mCherry (top panel) or GFP (bottom panel) (**E**). (**F**) RNAseq read counts of IFT-A, IFT-B, heterotrimer kinesin-II and BBSome genes in control and ANKRD55-transfected IMhu-M (24 h). Mean ± SEM from two independent experiments is presented. (**G**) qPCR analysis of the 8 IFT-B components found in the IMhu-M interactome in the 4 cell lines of the study. Mean ± SEM from three independent experiments is presented.

In order to determine the structural basis for the interaction of ANKRD55 with IFT-B components, we applied the visible immunoprecipitation (VIP) assay (35, 40). This assay facilitates direct visual assessment of binary and complex (one-to-many) protein interactions by fluorescence microscopy, and was used to identify the binding partner(s) of ANKRD55 in the IFT-B complex.

Lysates prepared from HEK293T cells co-expressing ANKRD55-EGFP and either all, multiple or single subunits from the IFT-B complex fused to mCherry were processed for immunoprecipitation with GST-anti-GFP nanobody prebound to glutathione-Sepharose beads. A red signal, implying retention of the mCherry-fused proteins on the agarose matrix via ANKRD55, was observed when ANKRD55-EGFP was co-expressed with either the complete IFT-B particle (Figure 7 B), the core, or IFT-B core-2, but not when co-expressed with the peripheral or IFT-B core-1 subcomplexes.

ANKRD55 co-expression with only four out of five IFT-B core-2 subunits resulted in a strong reduction of the red signal only when either IFT46 or IFT56, or to a lesser extent, IFT52 were absent (Figure 7 C). Figure 7 D shows that the smallest IFT-B core-2 substructure still capable of binding ANKRD55 was the IFT46 – IFT56 pair, given that the red signal was completely lost following co-expression of ANKRD55 with alternative pairs or individual core-2 subunits. Co-immunoprecipitation of mCherry-IFT46 and -IFT56 with ANKRD55-EGFP was confirmed by processing the immunoprecipitates for conventional immunoblotting (Figure 7 E).

The VIP data thus suggest that in IMhu-M cells, the eight identified IFT-B core proteins had been recruited to the ANKRD55 interactome, not through individual interaction with ANKRD55, but jointly with affinity-tagged ANKRD55 as a complex. We verified that transfection with the synthetic ANKRD55 RNA / nanoparticle combination used in this study had not artefactually induced IFT gene expression in the IMhu-M cell line. RNAseq data in Figure 7 F show that the 8 IFT genes as well as other genes involved in cilia biogenesis including additional genes of the IFT-B complex, and those of the IFT-A complex, heterotrimer kinesin-II group and BBSome, a protein complex that operates in primary cilia biogenesis, IFT and homeostasis, were, in fact, expressed at similar or somewhat lower levels in transfected compared to untransfected cells. Moreover, mRNAs for the eight IFT-B subunits identified in the IMhu-M interactome including the ANKRD55-binding IFT46 – IFT56 pair were expressed at similar levels in IMhu-A and SH-SY5Y compared to IMhu-M cells as measured by qPCR (Figure 7 G). These observations fail to explain the lack or reduced enrichment of IFT-B components in the ANKRD55 interactomes from the former cell lines.

### ANKRD55 Localizes to the Centrosome or Basal body but not to Primary Cilia

In contrast to astrocytes and neuronal cells that can form primary cilia, capacity of microglia to generate primary cilia is controversial (41, 42). The enrichment of an IFT-B-like – ANKRD55 complex in a microglia cell line, but not in the former cell lines, may be indicative for a ciliogenesis-independent function and localization of this complex. We analyzed native ANKRD55 co-localization with IFT-B components and other relevant markers of cilia, by means of confocal immunofluorescence microscopy in the untransfected IMhu-A, IMhu-M and SH-SY5Y cell lines.

This was done under conditions of serum starvation, which induces both cell cycle arrest and cilia formation, or under unstarved conditions, as indicated. First, we compared IF staining of two rabbit anti-human ANKRD55 Abs available as Prestige Antibodies® Powered by Atlas Antibodies, anti-ANKRD55 Ab1 and Ab2 (**Supplementary Table 1**) under normal cell culture conditions. These Abs were generated through immunization with discrete N- and C-terminal ANKRD55 antigen sequences. Both these Abs recognize proteins encoded by ANKRD55 transcripts 201 (69 kDa) and 203 (63.5 kDa) but only Ab2 recognizes transcript 202 isoform (37 kDa). In our earlier study (9), in SH-SY5Y cells, native 69 and 63 kDa ANKRD55 immunoreactive proteins were relatively enriched in the nuclear fractions, whereas a 37 kDa variant was more present in the cytosol and associated with membranous organelle fractions. In IF microscopy, Ab1 and Ab2 showed distinct staining patterns (Figure 8).

**FIGURE 8.**
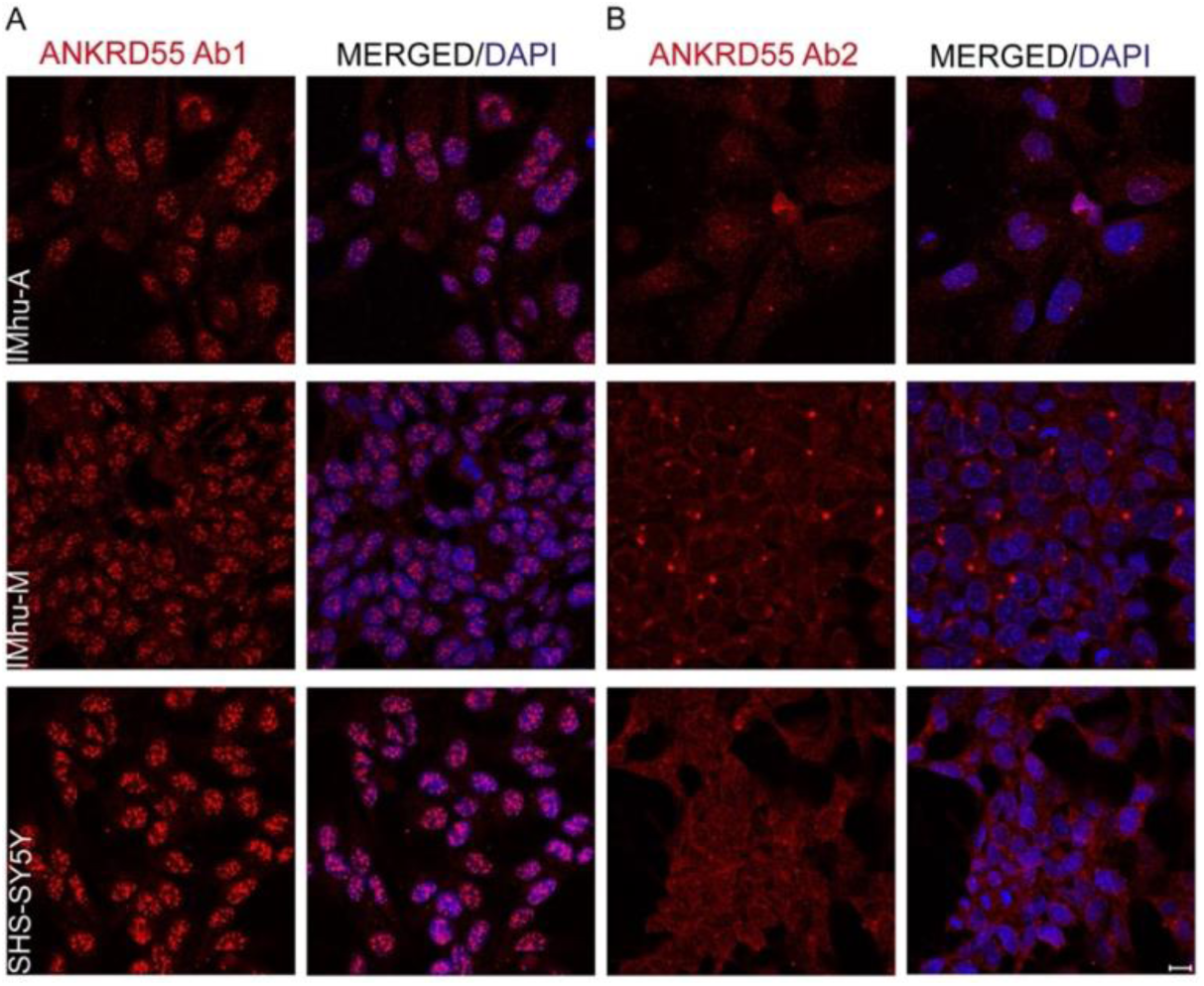
Differential immunodetection patterns of ANKRD55 by (A) Ab1 and (B) Ab2 in untransfected IMhu-A, IMhu-M and SH-SY5Y cell cultures. Scale bar: 10 μm.

In the three tested cell lines, Ab1-stained ANKRD55 was present in a pattern of bright speckles confined to the DAPI-stained nucleus with much weaker staining in the cytosol. Ab2 yielded a more diffuse granular staining pattern throughout both the nucleoplasm and cytosol, but also marked single intense spots adjacent to the nucleus, most clearly so in IMhu-M and IMhu-A cell lines. Ab2 was used for ensuing IF microscopy unless otherwise indicated. Cilia formation in IMhu-A, IMhu-M, SH-SY5Y cells was probed using ARL13B Ab, a commonly used marker of primary cilia (43). NIH-3T3 cells, a prevalent model for the study of cilia, were used as positive control (44). Abs against IFT46 and IFT56 compatible with the rabbit ANKRD55 Abs used in this study are commercially not available; therefore, we settled on use of a compatible IFT81 Ab, and we validated some key findings with a goat IFT74 Ab in combination with a rabbit IFT46 Ab (**Supplementary Table 1**). Both IFT74 and IFT81 are among the top ranked proteins identified in the IMhu-M ANKRD55 interactome (Figure 3 B). Following 24 h of serum deprivation, large ARL13B^+^ cilia were observed in IMhu-A cells, at a rate of about 1 cilium per cell, while also SH-SY5Y and NIH-3T3 cells formed cilia, which were smaller, and in SH-SY5Y cells less frequent (Figure 9 A). In contrast to the IFT-B component IFT81, which was visible in typical ciliary transport trains of these three cell lines (Figure 9 B), ANKRD55 was undetectable within the cilia of any cell line but was present, similar to IFT81, in bright spots adjacent to one of both tips of the cilia in IMhu-A and SH-SY5Y cell lines. ARL13B^+^ cilia were not observed in IMhu-M cells, even after prolonged serum deprivation (48 h), and in these cells ARL13B co-localized with either ANKRD55 or IFT81 in large bright spots in the cytosol. Via co-staining with an Ab for tubulin-γ1 (TUBG1), a protein associated with microtubule organizing centers (MTOC), these ANKRD55 – IFT81 co-localization areas in IMhu-M were identified as the centrosome (Figure 9 C). In ciliated cells, ANKRD55 was detected at the basal body, a protein structure also marked by TUBG1 that is found at the base of a cilium and that is derived from the mother centriole of the centrosome (Figure 9 D). Thus, ANKRD55 – IFT81 co-localization seem to occur at the mother centriole. We also tested co-localization of distinct IFT-B subunits in IMhu-M cells using a commercially available suitable selection of compatible antibodies. This showed that IFT81 co-localized with two tested IFT-B components, IFT46 and IFT74, mainly in the characteristic cytosolic areas of IMhu-M (Figure 9 E) identified as the centrosome. Similar to IFT81, IFT74 was detectable in cilia of IMhu-A (Figure 9 F); however, ANKRD55 remained undetectable in IMhu-A and SH-SY5Y cilia when using alternative ANKRD55 Ab1 (Figure 9 G).

**FIGURE 9.**
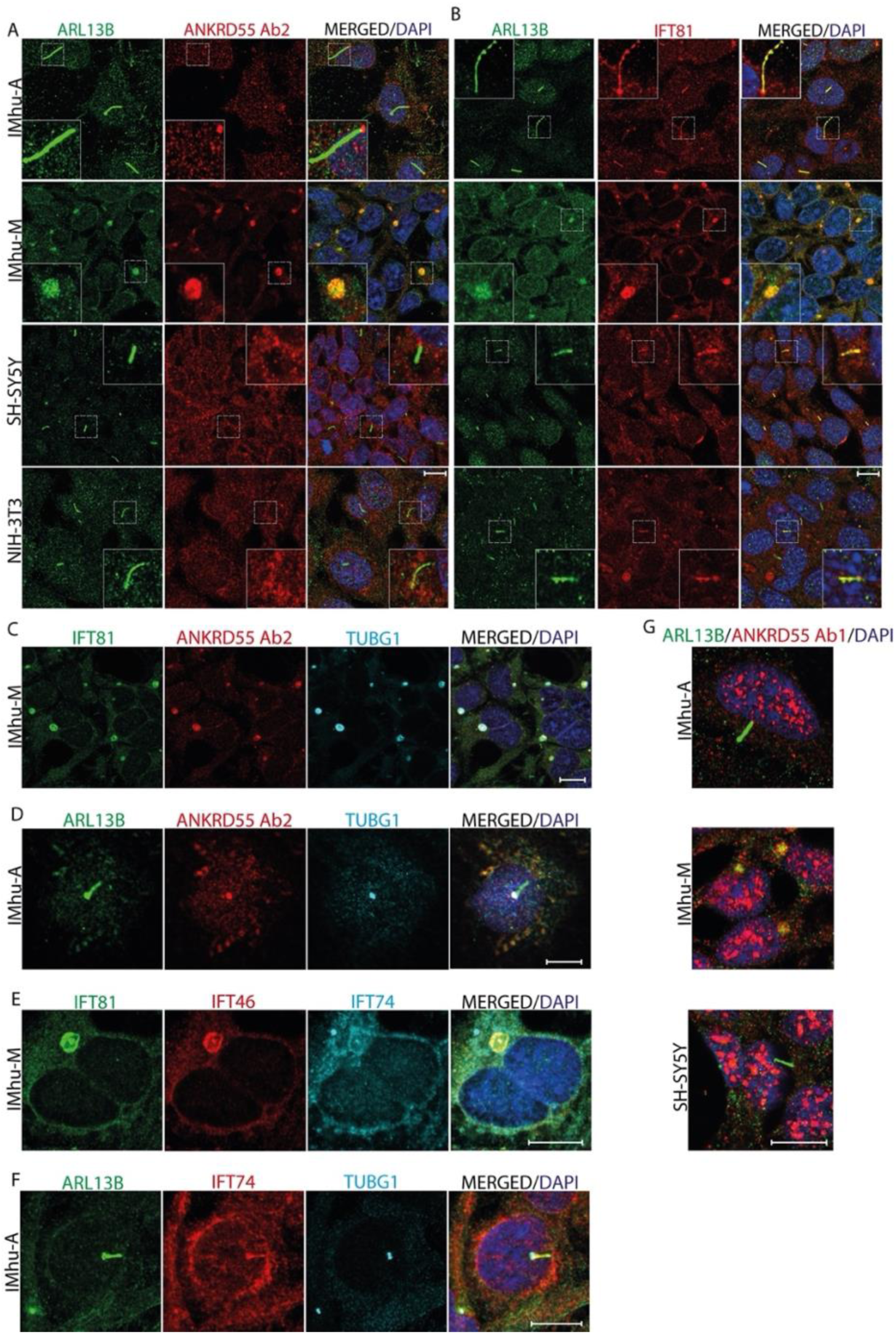
ANKRD55 is not detected in ARL13B^+^ primary cilia but localizes to the basal body in serum-starved astrocytic and neuronal cells, and to the centrosome in serum-starved non-ciliated microglial cells. Confocal microscopy co-localization patterns of ANKRD55 and cilium-complex proteins in serum-deprived IMhu-A, IMhu-M, SH-SY5Y and NIH-3T3 cells. Co-localization of (**A**) ARL13B and ANKRD55 (Ab2), (**B**) ARL13B and IFT81, (**C**) IFT81, ANKRD55 (Ab2) and TUBG1 in IMhu-M, (**D**) ARL13B, ANKRD55 (Ab2) and TUBG1 in IMhu-A, (**E**) IFT81, IFT46 and IFT74 in IMhu-M, (**F**) ARL13B, IFT74 and TUBG1 in IMhu-A, and (**G**) ARL13B and ANKRD55 (Ab1). Insets in (**A**) and (**B**) delineate magnified ciliar structures exhibiting a characteristic ARL13B expression in astrocytic, neuronal and fibroblast cell models. In the microglial cell model the insets show co-expression of ARL13B at centrosomal location together with ANKRD55 and IFT81 in the absence of any visible cilliary shape. Scale bar: 10 μm

Given support for a role for IFT subunits in vesicular trafficking in non-ciliated cells (45, 46), we asked whether ANKRD55 could be detected in association with specific vesicular structures. We performed a more quantitative approximation of the spatial relationship of immunofluorescence detection from triplets of proteins via antibody-linked 488, 594 and 647 fluorophores, as described in Materials and Methods. Specifically, we analysed the degree of spatial co-localization between ANKRD55 – IFT81 and markers distinguishing various types of vesicles in IMhu-M (not serum-starved). Figure 10 shows representative immunofluorescence images, digitalized outlines of signals and of triple merged areas (white fill), and Venn diagrams that provide a more quantitative estimate of overlapped areas. ANKRD55^+^ structures or ANKRD55-IFT81 overlapping areas showed only little spatial merging with markers of lysosome biogenesis and autophagy (LAMP-1 and LAMP-2), early endosomes (EEA1) and cis-Golgi stacks (GOLGA2). LC3^+^ autophagosomal structures appeared to congregate around the centrosome in this cell line, and showed stronger spatial overlap with ANKRD55-IFT81. We used bafilomycin A1, which is an inhibitor of vacuolar H^+^-ATPase and thereby inhibits the process of autophagy, to verify whether ANKRD55 was directly implied in autophagy. Bafilomycin A1 significantly increased LC3^+^ immunofluorescence, but appeared not to affect size, distribution or frequency of IFT81^+^, ANKRD55^+^ or TUBG1^+^ structures (**Supplementary Figure 3**). For comparison, we included in this analysis also TUBG1. Compared to the vesicular markers tested, spatial overlap was larger with TUBG1. The enrichment of ANKRD55 and IFT81 at the centrosome was validated with CEP43 (also known as FOP), a large centrosomal protein required for anchoring of microtubules at the centrosome (47). Spatial overlap of ANKRD55 – IFT81 with TUBG1 or LC3 in IMhu-A or SH-SY5Y was less pronounced than that seen in IMhu-M (**Supplementary Figure 4**).

**FIGURE 10.**
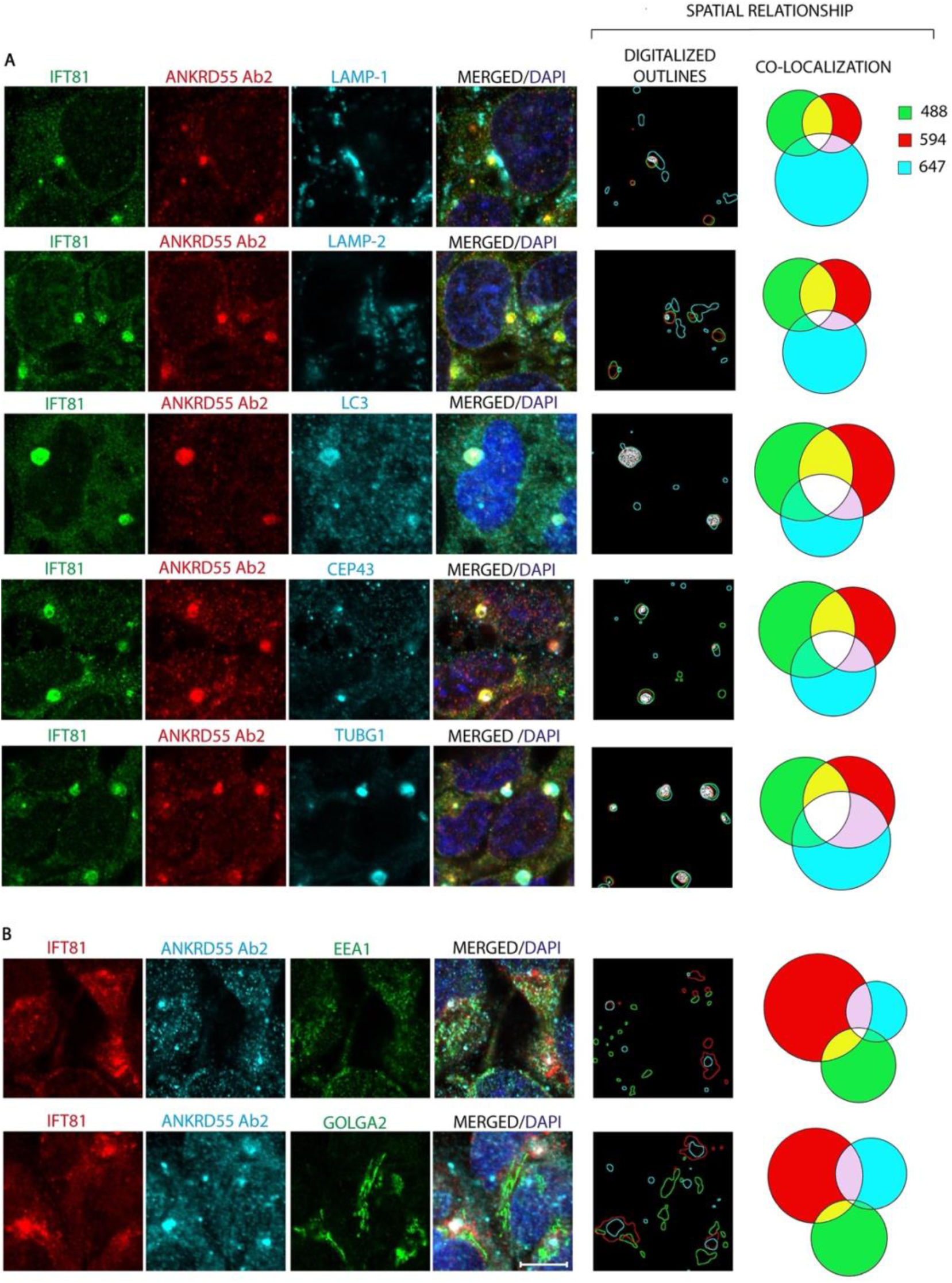
Immunofluorescence description of spatial relationship of IFT81 and ANKRD55 with vesicular markers in unstarved IMhu-M cells. Individual channel protein detection under the confocal microscope was outlined (see **Supplementary Figure 1**) and the defined areas were measured to generate corresponding Venn diagrams of protein co-localization. Abs against LAMP-1, LAMP-2, LC3, CEP43 and TUBG1 were labeled with Alexa647, and those against EEA1 and GOLGA2 with CoraLite488. Abs against IFT81 and ANKRD55 were labeled with (**A**) CoraLite488 and Alexa594, respectively, or (**B**) with CoraLite594 and Alexa647. In “Digitalized Outlines”, areas of selected immunostaining for each channel (488, 594, 647) were outlined with a custom-made Fiji macro and represented with the same color code. Triple immunodetection intersection areas are filled in white. Venn diagrams of the co-locatization of selected pixels were generated with a Python script. Scale bar: 10 µm.

We also assessed co-localization between ANKRD55 and 14-3-3 proteins. Using a pan-14-3-3 Ab, 14-3-3 proteins were found to be enriched at the centrosomal areas of IMhu-M, and no co-localization was observed with ANKRD55 outside the centrosome (**Supplementary Figure 5**).

Given the enrichment of mitochondrial membrane proteins in the SH-SY5Y interactome, we performed ANKRD55 co-staining with MitoTracker^TM^, a fluorescent dye that labels mitochondria in living cells. In none of the three tested cell lines did we observe any spatial co-localization. In a separate experiment, the mitochondrial protein ATP5PD, identified in two interactomes, did not co-localize noticeably with ANKRD55 (**Supplementary Figure 6**).

### Recapitulation of ANKRD55 – IFT81 Centrosomal Localization in Primary MoDC and MoMG

We asked whether the co-localization of ANKRD55 – IFT81 at the centrosome or with LC3 seen in the microglial cell line can be recapitulated in two primary CD14^+^ monocyte-derived cell models; MoDC, that express higher levels of ANKRD55 than monocytes (11), and MoMG, a primary model of authentic microglia exhibiting a phenotype and gene expression profile similar to human microglia (48). Differentiation of monocytes into MoDCs was performed in the presence of IL-4 / GM-CSF and was verified by enhanced expression of CD209 (Figure 11 A). MoMG were differentiated from monocytes in the presence of M-CSF, GM-CSF, NGF-β, CCL2 and IL-34, and were ascertained to express the microglia-specific marker P2RY12 (Figure 11 B) (31, 48). As in the IMhu-M cell line, ANKRD55 and IFT81 appeared to co-localize with bright spots positive for TUBG1 or LC3 in both MoDC and MoMG (Figure 11 C and **D**). Not al cells displayed co-localization of ANKRD55 – IFT81 with these spots. Co-localization of ANKRD55 – IFT81 with TUBG1 was seen more frequently than with LC3.

**FIGURE 11.**
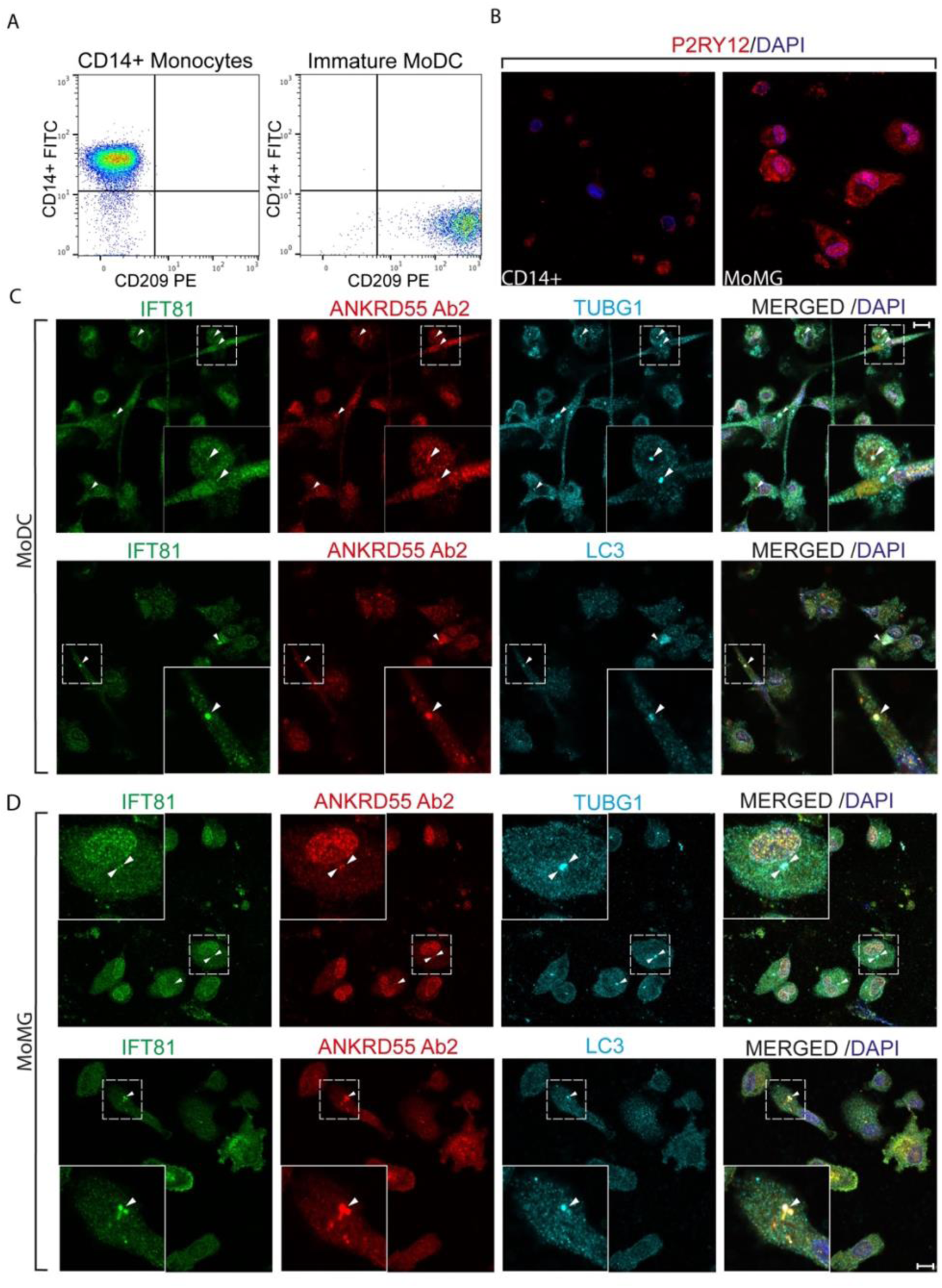
Co-localization of ANKRD55 and IFT81 with TUBG1 and LC3 in MoDC and MoMG by confocal microscopy. (**A**) Monocytes were differentiated into immature MoDCs by cultivation for 6 days in culture medium containing IL-4/GM-CSF. Differentiation was assessed by flow cytometry with CD14, a monocyte marker, and CD209, a marker for immature MoDC. (**B**) Monocytes were differentiated into MoMG in the presence of M-CSF, GM-CSF, NGF-β, CCL2 and IL-34 (31), for 8 days. Higher expression of microglia-specific marker P2RY12 in MoMG compared to monocytes was indicative for successful differentiation. (**C**) Co-localization by confocal microscopy of IFT81, ANKRD55 and TUBG1 or LC3 in MoDC. (**D**) Co-localization by confocal microscopy of IFT81, ANKRD55 and TUBG1 or LC3 in MoMG. Image insets show magnified stained areas. Scale bar: 10 µm.

## Discussion

In this work, ANKRD55 interactomes were determined from four distinct human cell types, astrocytic, microglial, neuroblastoma and monocytic, using nanoparticle-based transient transfection of synthetic ANKRD55 RNA and AP-MS. Although the majority of proteins detected in the interactomes were unique to each cell line, the results show a common core of ANKRD55-interacting proteins formed by three 14-3-3 proteins, 14-3-3η, 14-3-3β/α, and 14-3-3θ (encoded by *YWHAH*, *YWHAB*, and *YWHAQ* genes, respectively) shared by the four cell lines. Additional 14-3-3 isoforms were shared by the two glial and neuroblastoma cell lines, i.e. 14-3-3γ and 14-3-3ε (*YWHAG* and *YWHAE*, respectively), and both these isoforms have been identified also in published ANKRD55 interactomes from HEK293(T) cells (26, 27) and HCT116 cells (28). Based on NSAF (T–C) values, 14-3-3 proteins appeared to be the most abundant binding partners in the microglial ANKRD55 interactome, and were also highly enriched in the glial and neuroblastoma interactomes. 14-3-3 isoform binding could point to a range of diverse, well-documented cell signaling pathways in which ANKRD55 may be involved. However, since hundreds of functionally diverse 14-3-3 protein clients have been identified to date (49, 50), it is difficult to infer from this interaction precise biological processes in which ANKRD55 may participate. Bioinformatic analysis of the interactomes with exclusion of 14-3-3 proteins led to identification of the GO Component “intraciliary transport particle b”, uniquely in the microglia cell line. This term was the most significant of all annotations identified by STRING in the four individual non-14-3-3 category AB ANKRD55 interactomes. The protein network of this pathway comprised 8 IFT-B components, and using a VIP assay we show that ANKRD55 interacts specifically with the IFT46-IFT56 dimer of this complex. The source cell line IMhu-M, derived from primary human microglial cells, has been characterized in some detail and exhibits specific microglial markers and characteristics (51–53). In this study, this cell line did not form cilia under conditions of serum starvation.

Thus, we asked whether the biological context of the interaction of ANKRD55 with an IFT-B-like complex is pertinent to intraciliar transport. Substantial work has been done before by Drew and colleagues (22) who demonstrated that an ANKRD55-GFP fusion protein was actively transported within the cilia of multiciliated embryonal epithelial cells of the amphibia *X. laevis*. In our study, using confocal IF microscopy, we were unable to detect ANKRD55 in human neuronal and astrocytic primary cilia by means of Abs covering the known protein isoforms of ANKRD55, but we could identify it at the basal body. In the non-ciliated microglia cell line, ANKRD55 was enriched at the centrosomal area. A biological relationship exists between both organelles, given that the basal body emerges from the mother centriole of the centrosome (54). The variable intraciliar detectability of ANKRD55 could be related to differences in key features that set multicilia apart from monocilia (55, 56). The former are motile with sliding intraflagellar dynein arms on a specific axoneme microtubule structure, are post-mitotic and terminally differentiated. In contrast, the formation and resorption of primary cilia is dynamically regulated by the cell cycle (57). Multiciliated cells do not employ the semi-conservative centriole duplication program of cycling cells, i.e. centrosome consisting of mother and newly formed daughter centriole, and instead form numerous centrioles that are required for the growth of the motile cilia (58). In view of these discrepancies, a differential functional role for ANKRD55 in multicilia versus monocilia ciliogenesis can not be excluded.

However, our IF microscopy data argue against ANKRD55 being part of an intraciliarly transported IFT-B complex. Moreover, the comparative interactome proteomics, powered by the VIP assay used here, reveal that the IFT-B-like complex could be retrieved only from a cell line in which cilia could not be induced under conditions of serum starvation, but not from two cell lines, astrocytic and neuronal, that were capable of forming primary cilia. Thus, our study demonstrates the biological viability of an ANKRD55-IFT-B-like complex, and indicates that it may not primarily be associated with ciliogenesis and *de facto* intraciliar transport.

Multiple recent studies substantiate a role for IFT proteins in diverse cellular processes, outside the ciliary compartment. In T-lymphocytes, which lack cilia, IFT20, an IFT-B component of the peripheral subcomplex, translocates to the immune synapse, and induces formation of a complex with other IFT-B components (IFT20–IFT57–IFT88) and the TCR (59). In fact, it has been hypothesized that the immune synapse could represent the functional homolog of the primary cilium in non-ciliated hematopoietic cells, as both are characterized by polarized arrangement of centriole and Golgi, act as signaling platforms, and are sites of intense vesicular trafficking and targeted exocytosis (60). In macrophages, which do not assemble primary cilia either, depletion of IFT88 reduced the response to pro-inflammatory cues (61). Moreover, a series of studies have shown that IFT proteins contribute extensively to regulation of cell cycle progression or division via regulation of proper chromosome alignment, central spindle architecture, and astral microtubule formation and correct spindle orientation (reviewed in ref 62). An IFT46–IFT52–IFT70–IFT88 tetramer was shown to be necessary for efficient clustering of centrosomes during mitosis in cells harboring supernumerary centrosomes (63). This tetramer interacts directly with the mitotic kinesin motor HSET to ensure efficient centrosome clustering in mitosis.

Considering all ANKRD55 interactomes published to date (26–28)^4^, and including those presented in this study, IFT74 emerges as the protein most frequently identified, i.e. in 6 out of 9 independent interactome studies, followed by IFT46 (5 out of 9). IFT74 or the other IFTs identified in this study have been identified only very rarely or not at all in a repository of false-positive contaminants of the utilized FLAG-based affinity enrichment procedure (64). Since IFT74 does not seem to interact directly with ANKRD55, as shown in the VIP assay, it was likely identified in these studies as part of an IFT-B-type subcomplex that contains IFT46–IFT56. Bioplex (27, 28) identified five IFT-B components in the ANKRD55 interactome from HEK293T cells of which four, IFT46, IFT52, IFT70A and IFT74, but not IFT70B, are shared with the IMhu-M interactome of the present study. This suggests that the ANKRD55-associated IFT-B subcomplex may to a certain extent be variable, though inter-study technical variability of detection can also contribute. Though we did not perform an exhaustive analysis, IF co-localization studies did not yield strong indications that ANKR55 is associated with vesicular trafficking in IMhu-M cells. Quantitatively, IFT81-ANKRD55 co-localization was clearly concentrated at the centrosome, and also IFT46, IFT74 as well as 14-3-3 isoforms were enriched at the centrosome.

It remains to be determined if 14-3-3 proteins participate in a 3-way interaction with ANKRD55 and IFT-B components, or represent an unrelated process. The former is possible given that by binding to specific phosphorylated sites on target proteins, 14-3-3 proteins can force conformational changes or influence interactions between their targets and other molecules (65). Intrinsically disordered structures as seen in the C-terminal half of ANKRD55 are typically enriched in 14-3-3 clients (50).

In SH-SY5Y and THP-1 cells, mitochondrial membrane proteins were identified belonging to two important mitochondrial membrane complexes, the respiratory complex I (NDUF) – the first enzyme of the mitochondrial electron transport chain, and the ATP synthase complex (39). In addition, various members of two further, structurally distinct types of mitochondrial membrane proteins were identified, VDAC ion channels and SLC25 carriers (66). However, by confocal microscopy, native ANKRD55 did not appear to co-localize with mitochondria, casting doubt on a mitochondrial connection of importance. False-positive interactions, some of which can occur post-lysis, are documented in AP-MS studies (50, 67). In this case, excess levels of synthetic RNA-produced ANKRD55 far exceeding native levels could capture high amounts of natural 14-3-3 proteins, which are known to be the among the most abundant proteins in the cytoplasm (50). 14-3-3 proteins are also the most abundant proteins, as shown here, in the interactome of ANKRD55. Some of this 14-3-3, occurring in its active form as homodimers or 14-3-3 isoform heterodimers, may have been complexed with a different target through intermolecular bridge function of 14-3-3 dimers (68). These targets may be detectable in our set-up due to the high levels of ANKRD55 complexed with 14-3-3, coupled to non-biased sensitive mass spectrometry identification. In line with this hypothesis, multiple mitochondrial membrane proteins were identified in a cardiac 14-3-3 interactome study (69), and 14-3-3 is also well known as a regulator of mitochondrial ATP synthase (70).

In this study, we performed immunofluorescence staining assays with various antibodies to corroborate co-lozalization of ANKRD55 with selected highly ranked interactome proteins. The VIP assay allowed us to validate and deconstruct the binding of ANKRD55 with the IFT-B complex.

Additional proteins of interest are present in the interactomes that will need further scrutiny. Molecular biological experiments, e.g., bidirectional co-IP and Western blot, will need to be implemented in order to confirm the specificity of those. In further studies, proximity labeling via expression of a fusion protein of ANKRD55 and a proximity-based ligase, such as TurboID (71), could provide a more complete picture of ANKRD55’s dynamic interactome via the capture of weak and transient interactions.

In conclusion, we report the identification by AP-MS of an octamer IFT-B-like complex in the interactome of ANKRD55 from a human microglial cell line. By means of the VIP assay, we describe that the binding site for ANKRD55 in this complex is the IFT46-IFT56 pair. While we detected IFT74 and IFT81, two ANKRD55 interactors, in primary cilia of serum-starved astrocytic and neuroblastoma cell lines, we did not detect intraciliar ANKRD55. Enrichment of ANKRD55, as well as of IFT74 and IFT81, at centrosomal or basal body locations suggests the mother centriole as a site of their interaction. ANKRD55 may be a new player in specific aspects of centrosome biology including those demonstrated by earlier studies of IFT involvement (59–63).

## Author Contributions

JM, RTN, JDG, AA, AF, NUD and IA contributed to the acquisition and analysis of the experimental data. MA and FE performed mass spectrometry and proteomic data analysis. CL and OP provided expertise and antibodies for confocal microscopy. YK and KN performed the VIP assay and provided expertise. All authors contributed to the article and approved the submitted version.

## Acknowledgments

We are grateful to ISCIII, MINECO, GV and the Regional European Development (FEDER) for providing funding for this work.

## Data Availability Statement

The mass spectrometry data generated in this study was deposited in MassIVE database (MSV000093668). Link: https://massive.ucsd.edu/ProteoSAFe/static/massive.jsp

---

**Supplementary Table 1.**

| Antibody target | Host | Primary | Secondary | Fluorophore | Dilution | Supplier | Reference |
| --- | --- | --- | --- | --- | --- | --- | --- |
| ANKRD55 (Ab2) | Rabbit | ✓ |  | - | 1/200 | Sigma Aldrich | HPA051049 |
| ANKRD55 (Ab1) | Rabbit | ✓ |  | - | 1/200 | Sigma Aldrich | HPA061649 |
| ARL13B | Rabbit | ✓ |  | - | 1/500 | Protein Tech | 66739-1lg |
| ARL13B | Mouse | ✓ |  | - | 1/500 | Protein Tech | 17711-1-AP |
| ARL13B | Rabbit | ✓ |  | CoraLite® Plus 647 | 1/500 | Protein Tech | CL647-1771 |
| IFT81 | Rabbit | ✓ |  | CoraLite® Plus 488 | 1/500 | Protein Tech | CL488-11744 |
| IFT81 | Rabbit | ✓ |  | CoraLite® Plus 594 | 1/500 | Protein Tech | CL594-11744 |
| Gamma-1-Tubulin | Mouse | ✓ |  | - | 1/1000 | Thermo Fisher | MA1-19421 |
| IFT74 | Goat | ✓ |  | - | 1/200 | Thermo Fisher | PA1-9129 |
| IFT46 | Rabbit | ✓ |  | - | 1/200 | Sigma Aldrich | HPA057550 |
| LAMP1 | Mouse | ✓ |  | - | 1/200 | Hybridoma Bank | H4A3 |
| LAMP2 | Mouse | ✓ |  | - | 1/200 | Hybridoma Bank | H4B4 |
| LC3 | Mouse | ✓ |  | - | 1/200 | MBL | M152-3 |
| FGFROP | Mouse | ✓ |  | - | 1/1000 | Sigma Aldrich | H00011116-M01 |
| EEA1 | Mouse | ✓ |  | CoraLite® Plus 488 | 1/200 | Protein Tech | CL488-68065 |
| GOLGA2 | Mouse | ✓ |  | CoraLite® Plus 488 | 1/200 | Protein Tech | CL48811308 |
| P2RY12 | Rabbit | ✓ |  | - | 1/200 | Sigma Aldrich | HPA014518 |
| 14-3-3 Pan | Mouse | ✓ |  | - | 1/1000 | Thermo Fisher | MA5-12242 |
| ATP5H | Mouse | ✓ |  | - | 1/200 | Thermo Fisher | 459000 |
| Rabbit | Goat |  | ✓ | AlexaFluor 594 | 1/500 | Thermo Fisher | A-11072 |
| Rabbit | Donkey |  | ✓ | AlexaFluor 594 | 1/500 | ABCAM | ab150072 |
| Rabbit | Donkey |  | ✓ | AlexaFluor 647 | 1/500 | ABCAM | ab150075 |
| Goat | Donkey |  | ✓ | AlexaFluor 647 | 1/500 | ABCAM | Ab150139 |
| Mouse | Goat |  | ✓ | AlexaFluor 488 | 1/1000 | ABCAM | Ab150113 |
| Mouse | Goat |  | ✓ | Alexa Fluor 647 | 1/500 | Thermo Fisher | A32728 |
| Mouse | Goat |  | ✓ | Alexa fluor 594 | 1/500 | Thermo Fisher | A11005 |

## Supplementary Figures

**Supplementary Figure 1.**
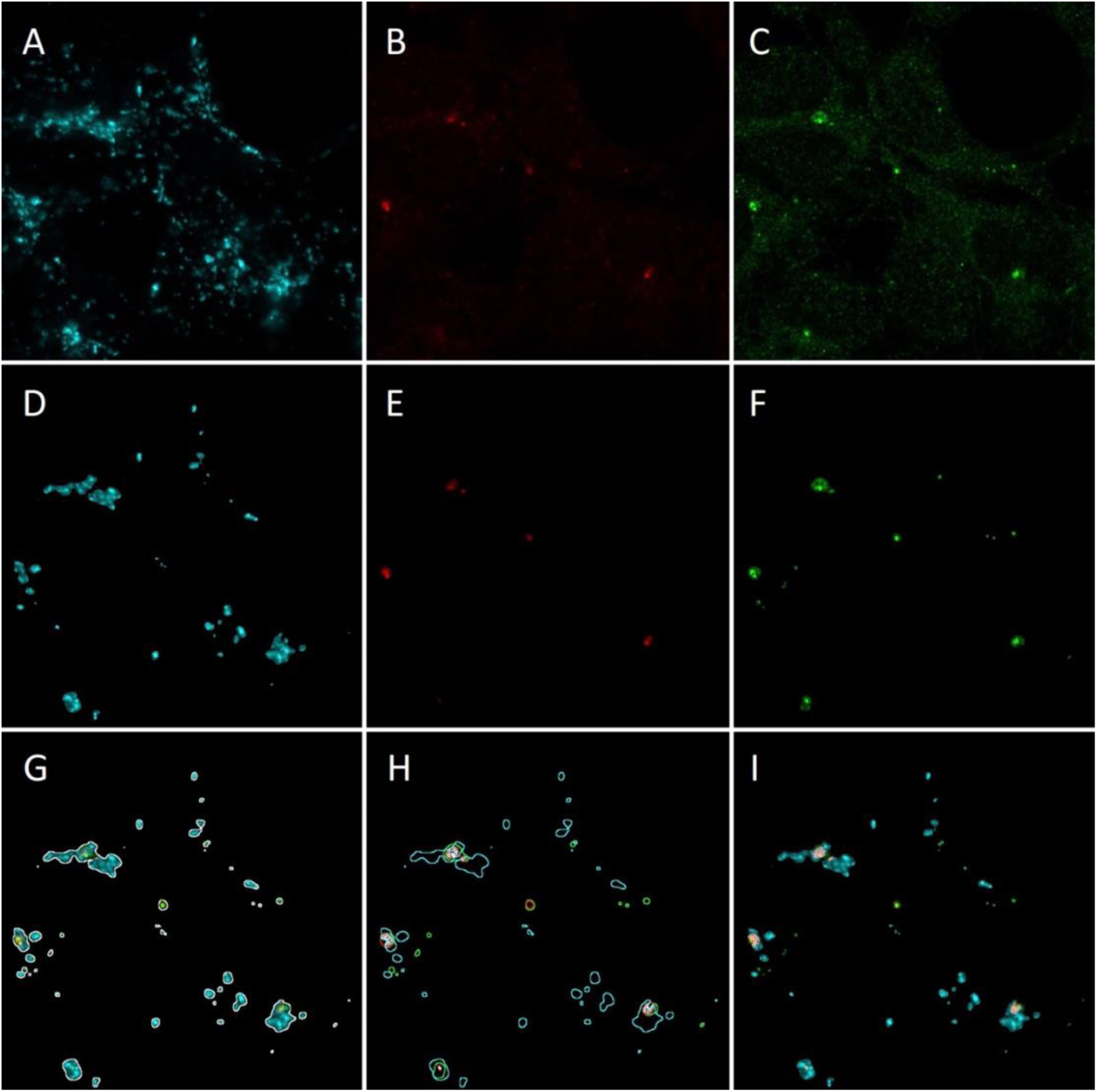
Image montage describing the analysis routine designed in Fiji software. Representative cropped area (830x830 pixels) of a multichannel-acquired image including fluorescent emissions at (**A**) 647 nm, (**B**) 594 nm and (**C**) 488 nm. (**D – F**) Manually intensity-thresholded images of corresponding wavelengths showing immunopositive selected regions in the cultured cells. (**G**) Fluorescent emissions of the three channels merged in a single RGB image with perimeters delineating total immunofluorescence area overlaid in white color. (**H**) Individual fluorescent emission perimeters in corresponding color (488nm-green, 594nm-red, 647nm-cyan) are overlapped on black background. Triple immunodetection intersection areas are filled in white. (**I**) Triple selected immunofluorescence-merged image with triple intersection areas overlaid in pink color.

**Supplementary Figure 2.**
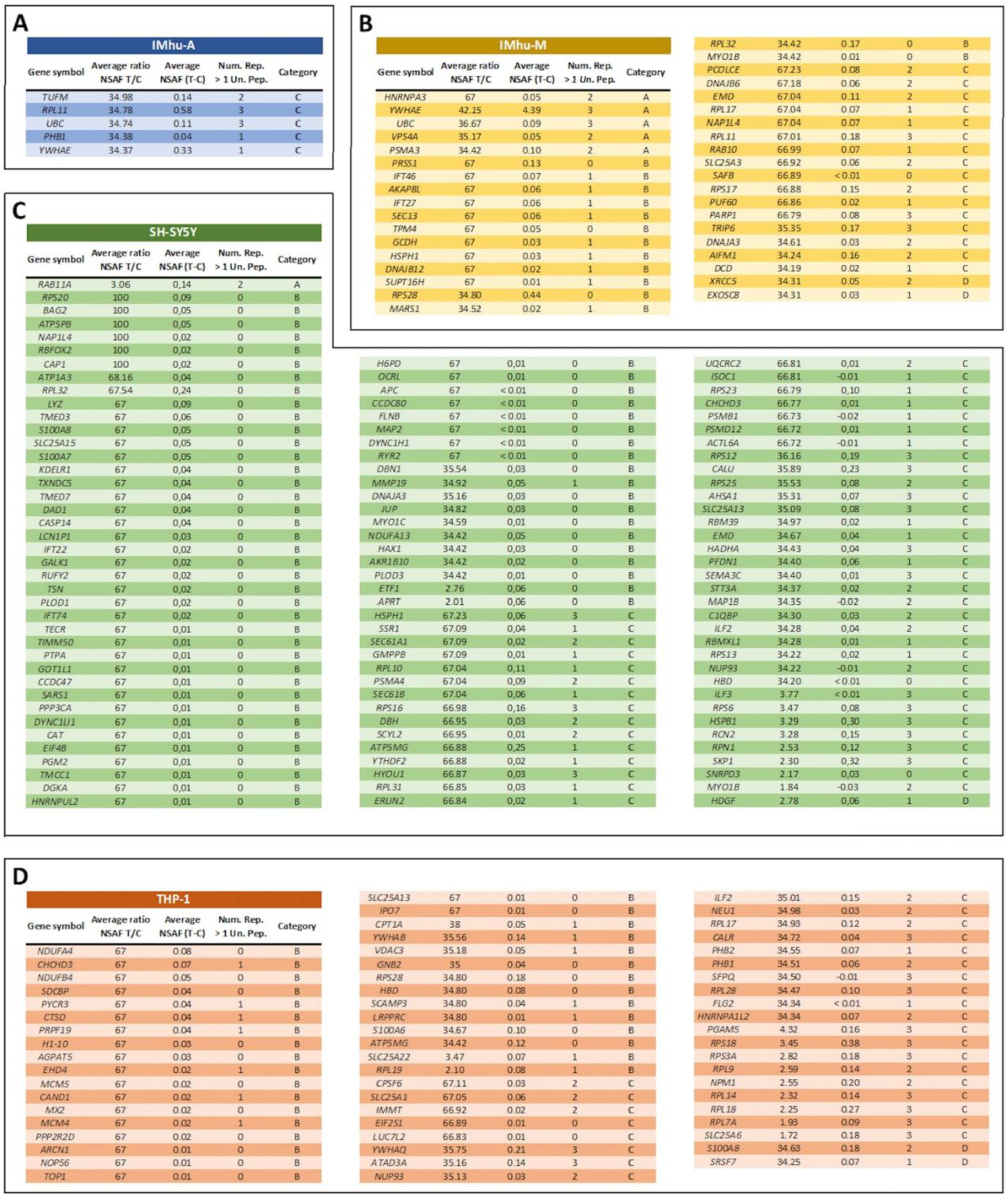
Remainder of proteins identified by FASP-based AP-MS in ANKRD55 interactomes of IMhu-A, IMhu-M, SH-SY5Y and THP-1 cells. Proteins are ranked according to the criteria of category A to D, and within each category according to degree of enrichment, i.e. from highest to lowest NSAF (T) / NSAF (C) ratio, averaged over three independent replicates. Proteins absent in the control and exclusively found in transfected cells in a single replicate are given a value of 100 for [Average ratio NSAF (T) / NSAF (C)]. Proteins absent in both control and transfected cells in a single replicate are given a value of 1. Number of replicates in which the protein was identified with more than one unique peptide is indicated (Num. Rep. > 1 Un. Pep.). Protein abundance is provided with NSAF (T – C).

**Supplementary Figure 3.**
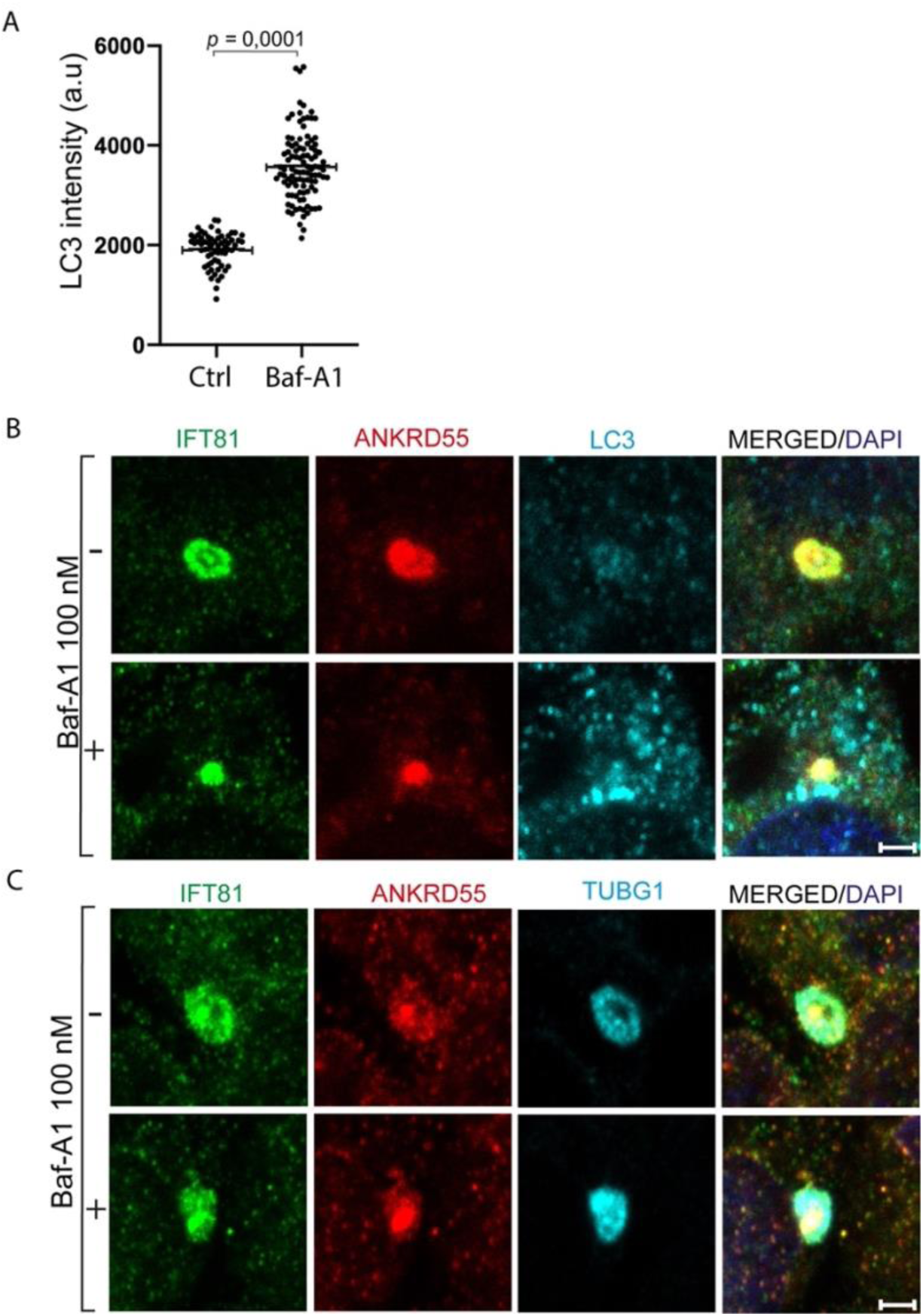
Analysis of autophagy inhibition with bafilomycin A1 treatment on appearance of IFT81^+^, ANKRD55^+^, TUBG1^+^ and LC3^+^ structures in confocal microscopy. IMhu-M cells were incubated with bafilomycin A1 (100 nM), or left untreated, for 24 h, immunostained with antibodies against IFT81, ANKRD55, and LC3 or TUBG1, and analyzed by confocal microscopy. (**A**) Scatter diagram of the average LC3 fluorescence intensity detected in centrosomal areas in cells cultured with and without bafilomycin A1 (Man Whitney test, *p* <0.0001). Confocal immunofluorescence triple detection of (**B**) LC3, IFT81 and ANKRD55, and (**C**) TUBG1, IFT81, ANKRD55, both in IMhu-M cells untreated and treated with bafilomycin A1. Scale bar: 10 µm.

**Supplementary Figure 4.**
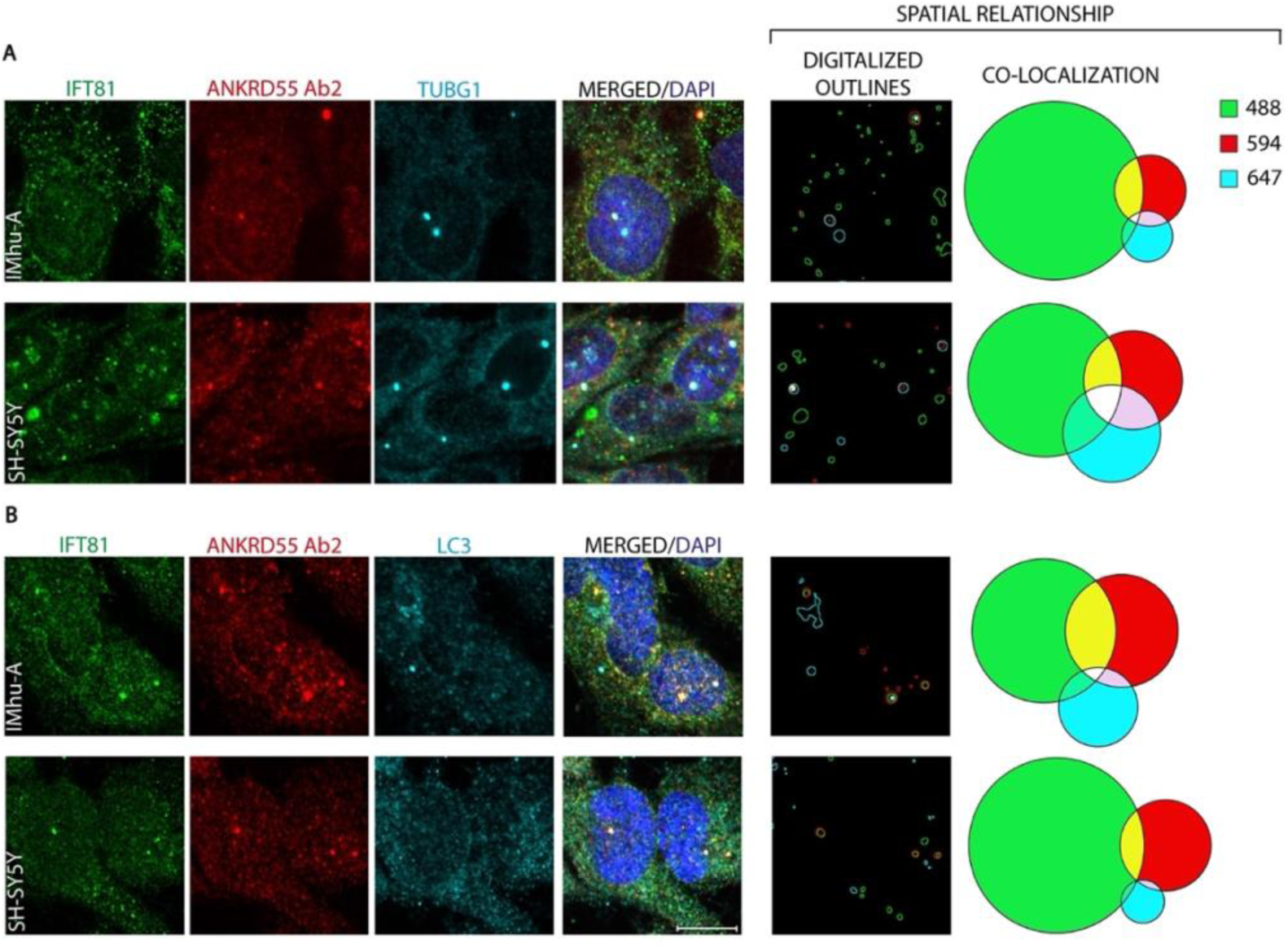
Immunofluorescence description of spatial relationship of IFT81 and ANKRD55 with (A) TUBG1 and (B) LC3 in IMhu-A and SH-SY5Y cells. Individual channel protein detection under the confocal microscope was outlined (see Supplementary Figure 1) and the defined areas were measured to generate corresponding Venn diagrams of protein co-localization. TUBG1 and LC3 were labeled with Alexa647. Abs against IFT81 and ANKRD55 were labeled with Coralite488 and Alexa594, respectively. In “Digitalized Outlines”, areas of selected immunostaining for each channel (488, 594, 647) were outlined with a custom-made Fiji macro and represented with the same color code. Venn diagrams of the co-localization of selected pixels were generated with a Python script. Scale bar: 10 µm.

**Supplementary Figure 5.**
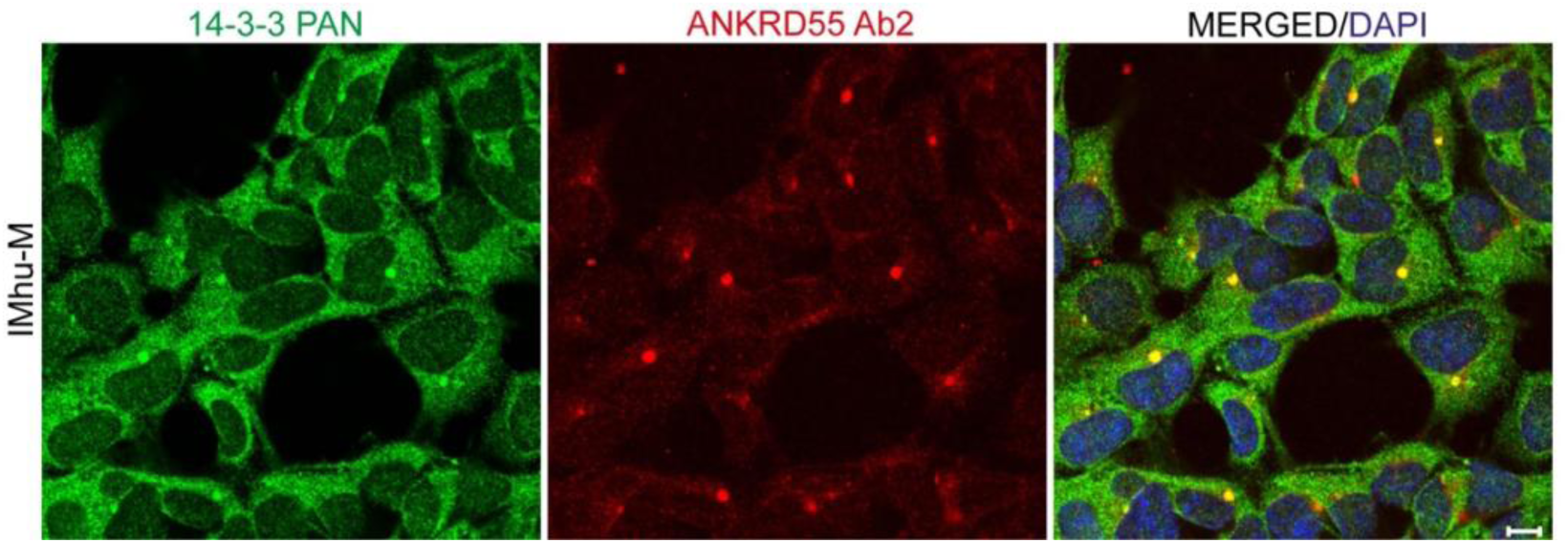
Confocal microscopy of ANKRD55 (Ab2), 14-3-3 (pan-14-3-3 Ab) and DAPI in IMhu-M cells. Scale bar: 10 µm.

**Supplementary Figure 6.**
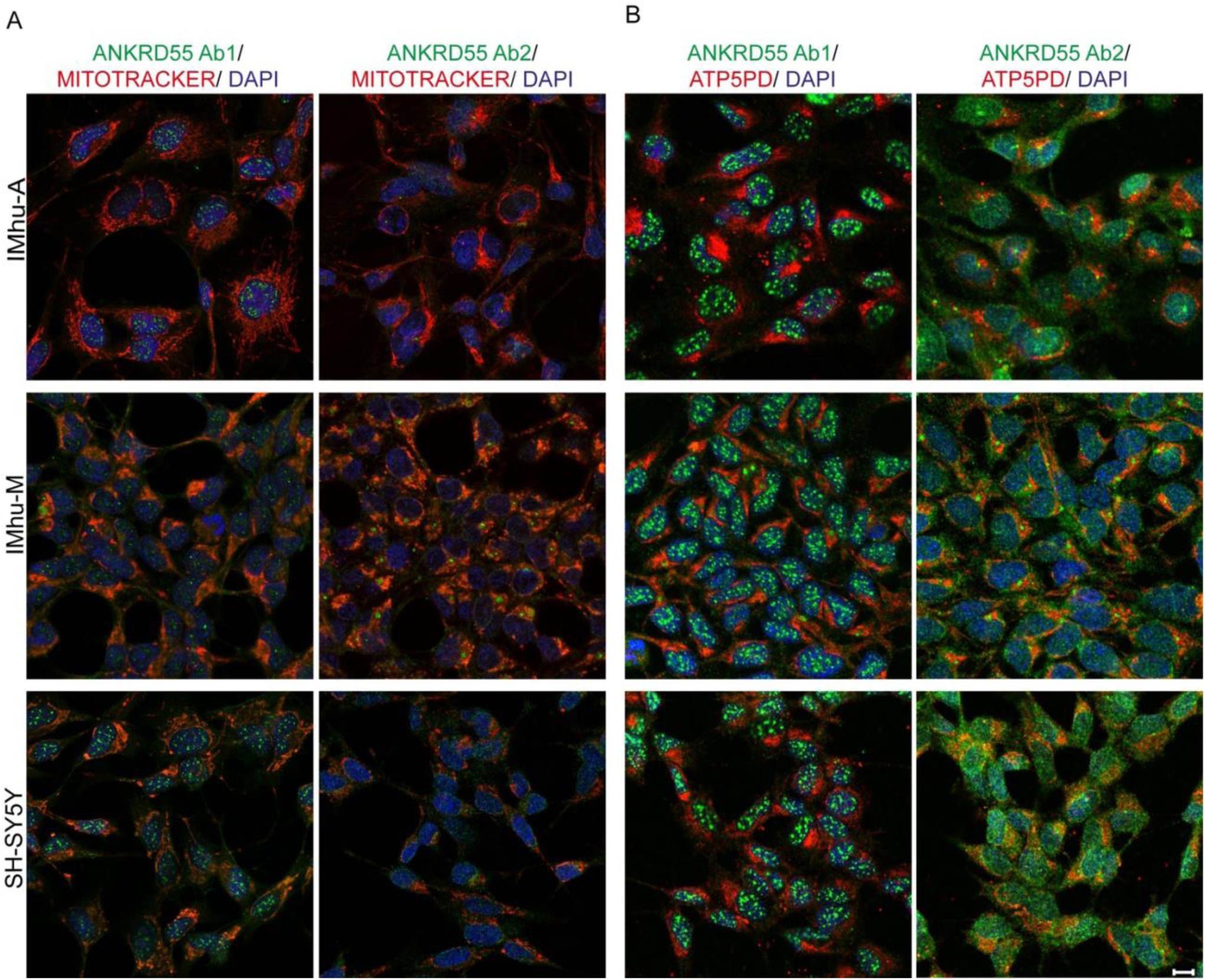
Confocal microscopy of IMhu-A, IMhu-M and SH-SY5Y cells stained with (A) MitoTracker^TM^, ANKRD55 (Ab1 and Ab2) and DAPI, and (B) ATP5PD, ANKRD55 (Ab1 and Ab2) and DAPI . Scale bar: 10 µm.

## Footnotes

1 https://gtexportal.org/home/gene/ANKRD55

2 https://www.proteinatlas.org/

3 https://bioplex.hms.harvard.edu/index.php

4 https://thebiogrid.org/122837/summary/homo-sapiens/ankrd55.html

